# Unearthing a fungal giant: *Dianjunaceae* fam. nov., a novel Paleocene lineage of *Xylariales* harbouring *Dianjunus rex* gen. et sp. nov.

**DOI:** 10.64898/2026.07.05.697275

**Authors:** Jing Song, Zhuyue Yan, Jesús Pérez-Moreno, Fengming Zhang, Shimei Yang, Tao Xie, Luofeng Su, Jianwei Liu, Yanliang Wang, Dong Liu, Zhenyan Yang, Chengmo Yang, Wei Liu, Xiaofei Shi, Shanping Wan, Ratchadawan Cheewangkoon, Dongqin Dai, Indunil C. Senanayake, Fuqiang Yu

## Abstract

During mycological surveys in Yunnan Province, China, specimens of a fungus producing massive, upright stromata up to 50 cm high and individually 2.2 Kg in weight were sampled. Through an integrative taxonomic approach combining detailed morphology, multilocus phylogeny (ITS, LSU, *RPB2*, *TUB2*), and phylogenomic analyses, this fungus is proposed as the new species *Dianjunus rex* gen. et sp. nov., the type of the new family *Dianjunaceae* (*Xylariales*). Phylogenetic analyses robustly place *Dianjunaceae* as a distinct sister clade to *Graphostromataceae*. Divergence time estimation dates the origin of this family to the early Paleocene (∼65 Mya), coinciding with the post-K-Pg extinction period, when an estimated 75% of all plant and animal species went extinct, and a significant ecological reorganization of life on earth happened. The stromata of *D. rex* represent the largest fructifications documented within the *Ascomycota*, significantly expanding the known morphological range of the *Xylariales*. The study provides a comprehensive description, including a nodulisporium-like anamorph with periconiella-like branching patterns, and discusses the taxon’s phylogenetic placement, and distinctive morphology. This discovery highlights the unexplored fungal diversity in East Asian forests.

## Introduction

The order *Xylariales* Nannf. comprises a morphologically diverse group of ascomycetous fungi, typically characterized by carbonaceous, darkly pigmented stromata. Taxa within this order occupy varied ecological niches, functioning as endophytes, pathogens, or saprobes (Senanayake et al. 2015). Saprobic members, often wood-associated, are key decomposers of dead plant materials, particularly lignified tissues, and contribute significantly to carbon cycling in forest ecosystems (Ma et al. 2022). Endophytic xylarialean species reside asymptomatically within healthy plant tissues, sometimes shifting to saprobic life mode upon host tissues senesce, while others may act as opportunistic plant pathogens under host stress (Cedeño–Sanchez et al. 2023).

*Xylariales* typically produce perithecial ascomata embedded within stromatic tissues. They are defined by unitunicate asci, frequently possessing a characteristic amyloid or inamyloid apical apparatus, and brown to dark ascospores that often exhibit a germ-slit (Stadler 2011). While traditional taxonomy has relied on stromatal morphology, perithecial arrangement, and spore characteristics, these traits are often subject to convergence. Contemporary systematics of *Xylariales* increasingly integrates multi-locus phylogenetic and genome-scale data, leading to substantial revisions of familial and generic boundaries (Wendt et al. 2018; Chen et al. 2023). Several studies indicated that anamorph morphology in *Xylariales* fungi served as a better indicator of phylogenetic relationships than teleomorph (Jaklitsch and Voglmayr 2012; Jaklitsch et al. 2016). The order currently encompasses 16 recognized families (Samarakoon et al. 2016) and allied lineages reflecting considerable lifestyle diversity. Beyond their ecological relevance, *Xylariales* are notable for their prolific production of secondary metabolites, including pigments and bioactive compounds, and their enzymatic capacity to degrade complex plant polymers. These attributes render them of significant interest for biotechnology, natural product discovery, and studies of plant–microbe interaction (Becker and Stadler 2021; Franco et al. 2025).

Stromatal morphology within the order varies, including both upright and penzigioid forms (Fournier et al. 2020). The order *Xylariales* encompasses few species capable of producing large stromata. A paramount example is *Engleromyces goetzei* P. Henn., which forms large, subglobose stromata on bamboo culms (Whalley et al. 2010). A documented specimen from Kenya comprised two lobes measuring 30 cm in diameter with a mass of approximately 3.2 kg, ranking among the most voluminous individual ascomata recorded in *Ascomycota* (Gibson and Kimeria 1962; Whalley et al. 2010). In contrast, species such as *Xylaria polymorpha* (Pers.) Grev. and *X. longipes* Nitschke produce clustered, clavate stromata where aggregations, though comprised of modest individual units (5–8 cm height), can exceed 20 cm in total breadth (Rogers 1979). *Squamotubera le-ratii* Henn., the only species of the genus reported with stromata around 7 cm height and 5 cm width (Hennings 1903; Rogers 1981).

During fungal diversity surveys in Yunnan Province, China, several specimens producing exceptionally large, upright stromata were collected from a tropical rainforest in Xishuangbanna Dai Autonomous Prefecture. Additional samples were obtained from temperate coniferous forests in Kunming and Lufeng Cities. Through integrative approaches, combining detailed morphological examination with multilocus phylogenetic reconstruction, taxonomic placement of all specimens were determined.

## Material and methods

### Sample collection, and morphological examination

Specimens were collected during the following periods and locations: August to September 2024 in Damaidi Village, Tuanjie Town, Luquan County, Kunming, China; July 2025 in Nanbang Village, Mengla County, Xishuangbanna Dai Autonomous Prefecture, Yunnan Province, China; and July 2025 in Yipinglang Town, Liupeng City, Yuzhu Yi Autonomous Prefecture, Yunnan Province, China. Detailed site data, including elevation, climatic conditions, and geographical features were recorded *in situ*. Photographs of fungal specimens in their natural habitat were taken using a Canon G15 camera (Canon Corporation, Tokyo, Japan). Collected materials were wrapped in water-proof foil and transported to the laboratory for subsequent analysis. Macroscopic characteristics were examined using an Olympus SZ61 stereomicroscope. Micromorphological structures, including ascospores, asci and ascomata were photographed using a Canon 700D digital camera mounted on a compound microscope. Ascospores were cultured on Potato Dextrose Agar (PDA), and pure cultures were obtained following the methodology of Senanayake et al. (2020a). Fungal hyphae were grown on Malt extract agar (MEA), Oat Meal Agar (OMA) and PDA to examine the colony morphology. All microscopic measurements were performed using the Tarosoft image Framework (v. 0.9.0.7). Image composites were processed and adjusted with Adobe Photoshop CS6 software (version 10.0, Adobe Systems, USA). Voucher specimens were deposited at the fungarium of the Kunming Institute of Botany Academia of Sciences (HKAS). Index Fungorum numbers (http://www.indexfungorum.org) were registered for all novel taxa.

### DNA extraction, PCR amplification, and sequencing

Genomic DNA was extracted from the fresh stromata tissue using the Biospin DNA Extraction Kit (Bioer Technology Co. Ltd., Hangzhou, China) according to the manufactureŕs protocol. Extracted DNA was stored at -20 °C prior to amplification. Polymerase chain reaction (PCR) amplification and sequencing were performed for three genetic loci: the partial 28S Large Subunit ribosomal DNA (LSU) using primers LR0R and LR5 (Vilgalys and Hester 1990); the complete Internal Transcribed Spacer (ITS) region using primers ITS1 and ITS4 (White et al. 1990); the beta-tubulin (*TUB2*) gene using primers T1 and T22 (O’Donnell and Cigelnik 1997); and the RNA polymerase II subunit 2 (RPB2) gene using primers fRPB2-5F and fRPB2-7cR (Liu et al. 1999). PCR reactions were conducted following the protocol described in Senanayake et al. (2020b). The total volume of the PCR reaction was 25 µL, containing 1 µL of DNA template, 1 µL of each forward and reverse primer, 12.5 µL of 2 × PCR Master Mix, and 9.5 µL of sterile double-distilled water (ddH_2_O). Amplification was performed for 35 cycles under the termocycling conditions detailed in Table 1. PCR products were visualized via electrophoresis on 1% agarose gels stained with ethidium bromide. Products were purified and sequenced commercially (Sunbiotech Company, Beijing, China). Raw sequences were assembled, edited and quality-checking using DNASTAR Lasergene v. 7.1. Newly generated sequences were deposited in the NCBI GenBank database; corresponding accession numbers are provided in Table 2.

**Table 1.** Primers and optimized PCR protocols used in this study.

| Locus | PCR protocols (Annealing temperature in bold) |
| --- | --- |
| LSU | (94 °C: 1 min, **56 °C**: 50s, 72 °C: 1 min) × 35 cycles |
| ITS | (94 : 50s, **56 °C**: 50s, 72 °C: 1 min) × 35 cycles |
| *TUB2* | (94 °C: 30s, **57 °C**: 50s, 72 °C: 1 min) × 35 cycles |
| *RPB2* | (95 °C: 30s, **55 °C**: 50s, 72 °C: 1 min) × 35 cycles |

**Table 2.** Details of the isolates used in the phylogenetic tree. Newly generated sequences are bold.

| Species | Culture accession number | ITS | LSU | *RPB2* | *TUB2* | References |
| --- | --- | --- | --- | --- | --- | --- |
| *Acrocordiella omanensis* | SQUCC 15091<sup>T</sup> | MG584568 | MG584570 | – | – | Maharachchikumbura et al. (2018) |
| *Acrocordiella yunnanensis* | HKAS 111922<sup>T</sup> | MW424507 | MW424505 | – | – | Dissanayake et al. (2021) |
| *Alloanthostomella rubicola* | MFLUCC 16-0479 | KX533455 | KX533456 | – | – | Daranagama et al. (2015) |
| *Allocryptovalsa cryptovalsoidea* | HVFIG02 | HQ692573 | – | – | HQ692524 | Trouillas et al. (2011) |
| *Allocryptovalsa elaeidis* | MFLUCC 15-0707<sup>T</sup> | MN308410 | MN308401 | – | MN340296 | Konta et al. (2020) |
| *Allocryptovalsa microtheca* | MFLUCC 18-1434 | PV617424 | PV617572 | PV902086 | PV901980 | Dissanayake et al. (2025) |
| *Allocryptovalsa polyspora* | MFLUCC 17-0364<sup>T</sup> | MF959500 | MF959503 | – | MG334556 | Senwanna et al. (2017) |
| *Allocryptovalsa rabenhorstii* | WA08CB | HQ692619 | – | – | HQ692523 | Senwanna et al. (2017) |
| *Allocryptovalsa truncate* | PUFNI17639<sup>T</sup> | MK990279 | MK981538 | – | – | Hyde et al. (2020) |
| *Allodiatrype arengae* | MFLUCC 15-0713<sup>T</sup> | MN308411 | MN308402 | MN542886 | MN340297 | Konta et al. (2020) |
| *Alloeutypa flavovirens* | CHUNI6 | KR092798 | KR092774 | – | – | Senanayake et al. (2015) |
| *Amphibambusa bambusicola* | MFLUCC 11-0617<sup>T</sup> | KP744433 | KP744474 | – | – | Liu et al. (2015) |
| *Amphirosellinia nigrospora* | HAST 91092308<sup>T</sup> | GU322457 | – | GQ848340 | GQ495951 | Hsieh et al. (2010) |
| *Annulohypoxylon terebratum* | CBS 119137<sup>T</sup> | DQ631943 | DQ840069 | DQ631954 | DQ840097 | Fournier et al. (2010) |
| *Annulohypoxylon truncatum* | CBS 140778<sup>T</sup> | KY610419 | KY610419 | KY624277 | KX376352 | Wendt et al. (2018) |
| *Anthostoma decipiens* | CD | KC774565 | KC774565 | – | – | Jaklitsch et al. (2014) |
| *Anthostomella obesa* | MFLUCC 14-0171<sup>T</sup> | KP297405 | KP340546 | KP340533 | KP406616 | Daranagama et al. (2015) |
| *Anthostomelloides krabiensis* | MFLUCC 15-0678 | KX305927 | KX305928 | KX305929 | – | Tibpromma et al. (2017) |
| *Arecophila miscanthi* | FU31025 | MK503821 | MK503827 | – | – | Karunarathna et al. (2021) |
| *Ascotricha chartarum* | CBS 902.69 | MH859477 | MH871257 | – | – | Vu et al. (2019) |
| *Ascotricha erinacea* | CBS 535.73 | KF893285 | MH872482 | – | KF893272 | Cheng et al. (2015) |
| *Ascotricha rugispora* | NBRC 33167<sup>T</sup> | LC146763 | LC146763 | – | – | Okane et al. (2001) |
| *Astrocystis mirabilis* | HAST 94070803 | GU322448 | – | GQ844835 | GQ495941 | Hsieh et al. (2010) |
| *Barrmaelia macrospora* | CBS 142768<sup>T</sup> | KC774566 | KC774566 | MF488995 | MF489014 | Jaklitsch et al. (2014) |
| *Barrmaelia rappazii* | CBS 142771<sup>T</sup> | MF488989 | MF488989 | MF488998 | MF489017 | Voglmayr et al. (2018) |
| *Barrmaelia rhamnicola* | CBS 142772<sup>T</sup> | MF488990 | MF488990 | MF488999 | MF489018 | Voglmayr et al. (2018) |
| *Biscogniauxia anceps* | YMJ 123 | EF026132 | – | JX507777 | AY951671 | Hsieh et al. (2010) |
| *Biscogniauxia capnodes* | YMJ 138 | EF026131 | – | JX507779 | AY951675 | Hsieh et al. (2010) |
| *Biscogniauxia mangiferae* | MFLU 18-0827<sup>T</sup> | MN337232 | MN336236 | MN366247 | MN509782 | Hyde et al. (2020) |
| *Biscogniauxia nummularia* | MUCL 51395<sup>T</sup> | KY610382 | KY610427 | KY624236 | KX271241 | Wendt et al. (2018) |
| *Brunneiperidium gracilentum* | MFLUCC 14-0011<sup>T</sup> | KP297400 | KP340542 | KP340528 | KP406611 | Daranagama et al. (2015) |
| *Cainia graminis* | MFLUCC 15-0540 | KR092793 | KR092781 | – | – | Senanayake et al. (2015) |
| *Calceomyces lacunosus* | CBS 633.88<sup>T</sup> | KY610397 | KY610476 | KY624293 | KX271265 | Wendt et al. (2018) |
| *Camillea obularia* | ATCC 28093 | KY610384 | KY610429 | KY624238 | KX271243 | Wendt et al. (2018) |
| *Castellaniomyces rosae* | MFLUCC 15-0536<sup>T</sup> | MF614127 | MF614130 | – | – | Senanayake et al. (2015) |
| *Circinotrichum maculiforme* | CBS 122758 | KR611875 | KR611896 | – | – | Crous et al. (2015) |
| *Clypeosphaeria mamillana* | CBS140735<sup>T</sup> | KT949897 | KT949897 | MF489001 | MH704637 | Jaklitsch et al. (2016) |
| *Clypeosphaeria mamillana* | WU 33599 | KT949898 | KT949898 | – | – | Jaklitsch et al. (2016) |
| *Clypeosphaeria mamillana* | WU 36850 | KT949899 | KT949899 | – | – | Jaklitsch et al. (2016) |
| *Clypeosphaeria perfidiosa* | CBS 142773 | MF488993 | – | MF489003 | MF489021 | Vu et al. (2019) |
| *Collodiscula japonica* | CBS 124266 | JF440974 | MH874889 | KY624273 | – | Jaklitsch and Voglmayr (2012) |
| *Coniocessia anandra* | CBS 125766<sup>T</sup> | MH863747 | MH875215 | – | – | Vu et al. (2019) |
| *Coniocessia cruciformis* | CBS 126674 | – | MH875663 | – | – | Vu et al. (2019) |
| *Coniocessia maxima* | CBS 593.74<sup>T</sup> | GU553332 | MH878275 | – | – | Vu et al. (2019) |
| *Coniocessia nodulisporioides* | CBS 281.77<sup>T</sup> | MH861061 | MH872831 | – | – | Vu et al. (2019) |
| *Coniolariaella gamsii* | Co27 | GU553325 | GU553329 | – | – | Vu et al. (2019) |
| *Creosphaeria sassafras* | STMA 14087 | KY610411 | KY610468 | KY624265 | KX271258 | Wendt et al. (2018) |
| *Cryptosphaeria eunomia* | CBS216.87<sup>T</sup> | KT425230 | KT425296 | – | – | Trouillas et al. (2015) |
| *Cryptosphaeria subcutanea* | CBS 240.87<sup>T</sup> | KT425232 | KT425297 | – | – | Trouillas et al. (2015) |
| *Cryptostroma corticale* | CBS 216.52 | HG934110 | MH868529 | HG934116 | HG934102 | Vu et al. (2019) |
| *Cryptovalsa ampelina* | ATCC MYA-4399 | FJ430591 | FJ430582 | – | – | Trouillas et al. (2010) |
| *Daldinia concentrica* | CBS 113277 | AY616683 | KY610434 | KY624243 | KC977274 | Triebel et al. (2005) |
| *Daldinia loculatoidea* | CBS 113279<sup>T</sup> | AF176982 | KY610438 | KY624247 | KX271246 | Triebel et al. (2005) |
| *Daldinia* sp. | ATCC 46302 | KY610389 | KY610443 | KY624253 | KX271248 | Wendt et al. (2018) |
| *Delonicicola siamense* | MFLUCC 15-0670<sup>T</sup> | MF167586 | MF158345 | MF158346 | – | Perera et al. (2017) |
| *Dematophora bunodes* | CBS 124028 | MN984619 | MN984625 | – | MN987245 | Wittstein et al. (2020) |
| *Dematophora necatrix* | 89062904 | EF026117 | – | – | EF025603 | Hsieh et al. (2010) |
| *Dematophora necatrix* | CBS 349.36 | AY909001 | KF719204 | KY624275 | KY624310 | Pelaez et al. (2008) |
| *Diabolocovidia claustris* | CPC37593<sup>T</sup> | MT373367 | MT373350 | – | – | Crous et al. (2020) |
| ***Dianjunus rex*** | **HKAS148941<sup>T</sup>** | **PX777991** | **PX777996** | **PX987258** | **PX930195** | **This study** |
| ***Dianjunus rex*** | **HKAS148942** | **PX777992** | **PX777997** | **PX987259** | **PX930196** | **This study** |
| ***Dianjunus rex*** | **HKAS150823** | **PX777993** | **PX777998** | **PX987260** | – | **This study** |
| ***Dianjunus rex*** | **HKAS150824** | **PX777994** | **PX777999** | **PX987261** | **PX930197** | **This study** |
| ***Dianjunus rex*** | **HKAS151626** | **PX777995** | **PX778000** | **PX987262** | – | **This study** |
| *Diatrype bullata* | UCDDCH400 | DQ006946 | – | – | DQ007002 | Rolshausen et al. (2006) |
| *Diatrype disciformis* | AFTOL-ID 927 | DQ470964 | U47829 | DQ470915 | – | Spatafora et al. (2006) |
| *Diatrype lijiangensis* | MFLU 19-0717<sup>T</sup> | MK852582 | MK810546 | – | MK852583 | Thiyagaraja et al. (2019) |
| *Diatrype virescens* | CBS 128344 | MH864890 | MH876339 | – | – | Vu et al. (2019) |
| *Diatrypella favacea* | CBS 198.49 | MH856491 | MH868027 | – | – | Vu et al. (2019) |
| *Diatrypella heveae* | MFLUCC 15-0274<sup>T</sup> | MN308418 | MN308409 | MN542892 | MN340301 | Vu et al. (2019) |
| *Diatrypella tectonae* | MFLUCC 12-0172b<sup>T</sup> | KY283085 | KY283080 | – | – | Shang et al. (2017) |
| *Didymobotryum rigidum* | JCM 8837<sup>T</sup> | LC228650 | LC228707 | – | – | Schoch et al. (2012) |
| *Discoxylaria myrmecophila* | 169 | GU322433 | – | GQ844819 | GQ487710 | Hsieh et al. (2010) |
| *Durotheca crateriformis* | GMBC 0205<sup>T</sup> | MH645426 | MH645425 | MH645427 | MH049441 | De Long et al. (2019) |
| *Emarcea castanopsidicola* | CBS 117105<sup>T</sup> | AY603496 | MK762717 | MK791285 | MK776962 | Duong et al. (2004) |
| *Endocalyx cinctus* | JCM 7946 | LC228648 | LC228704 | – | – | Schoch et al. (2012) |
| *Endocalyx metroxyli* | MFLUCC 15-0723A<sup>T</sup> | MT929162 | MT929313 | – | – | Thitla et al. (2025) |
| *Engleromyces sinensis* | BJTC 200802<sup>ET</sup> | MZ622704 | MZ622701 | MZ622185 | – | Gao et al. (2023) |
| *Engleromyces sinensis* | BJTC 200801 | MZ622703 | MZ622700 | MZ622184 | MZ622187 | Gao et al. (2023) |
| *Entalbostroma erumpens* | ICMP 21152<sup>T</sup> | KX258206 | – | KX258204 | KX258205 | Johnston et al. (2016) |
| *Entosordaria quercina* | CBS 142774<sup>T</sup> | MF488994 | MF488994 | MF489004 | MF489022 | Voglmayr et al. (2018) |
| *Eutypa lata* | DSC100 | HM164729 | – | HM164797 | HM164763 | Trouillas and Gubler (2010) |
| *Eutypella quaternata* | MFLU 15-2605 | MT185553 | MT183518 | MT432247 | – | Li et al. (2020) |
| *Fasciatispora arengae* | MFLUCC 15-0326a<sup>T</sup> | MK120275 | MK120300 | MK890794 | MK890793 | Doilom et al. (2018) |
| *Fasciatispora cocoas* | MFLUCC 18-1445<sup>T</sup> | MN482680 | MN482675 | MN481517 | MN505154 | Hyde et al. (2020) |
| *Fasciatispora nypae* | MFLUCC 11-0382<sup>T</sup> | NR_185352 | NG_067532 | – | – | Liu et al. (2015) |
| *Fusiformiascus pulvinatus* | H048 | FR715523 | FR715523 | – | FR715495 | Pazoutova et al. (2012) |
| *Graphostroma platystomum* | CBS 270.87<sup>T</sup> | JX658535 | DQ836906 | KY624296 | HG934108 | Stadler et al. (2014) |
| *Guayaquilina cubensis* | MUCL 39017 | KC775733 | KC775708 | – | – | Becerra-Hernández et al. (2016) |
| *Gyrothrix inops* | BE108 | KC775746 | KC775721 | – | – | Becerra-Hernández et al. (2016) |
| *Gyrothrix ramose* | MUCL 54061 | KC775747 | KC775722 | – | – | Becerra-Hernández et al. (2016) |
| *Halocryptovalsa salicorniae* | MFLUCC 15-0185<sup>T</sup> | MH304410 | MH305304 | – | MH370274 | Dayarathne et al. (2020) |
| *Halodiatrype avicenniae* | MFLUCC 15-0948 | MH304414 | MH305308 | – | MH370278 | Dayarathne et al. (2020) |
| *Halorosellinia krabiensis* | MFLU 17-2596<sup>T</sup> | MN047119 | MN017883 | – | MN431493 | Dayarathne et al. (2020) |
| *Hansfordia pruni* | CBS 194.56<sup>T</sup> | MK442585 | MH869122 | KU684307 | – | Crous et al. (2019) |
| *Hansfordia pulvinata* | CBS 144422 | MK442587 | MK442527 | – | – | Crous et al. (2019) |
| *Hansfordia pulvinata* | CBS 254.59 | KF893288 | MH869394 | – | – | Crous et al. (2019) |
| *Haploanthostomella elaeidis* | MFLU 20-0522<sup>T</sup> | MT929161 | MT929312 | – | – | Thitla et al. (2025) |
| *Hypocopra anomala* | TTI-000339 | – | MT903245 | MT901033 | MT901030 | Becker et al. (2020) |
| *Hypocopra dolichopoda* | TTI-000310 | – | MT903247 | MT901035 | – | Becker et al. (2020) |
| *Hypocopra rostrata* | NRRL 66178 | KM067909 | KM067909 | – | – | Jayanetti et al. (2014) |
| *Hypomontagnella monticulosa* | MFLUCC 18-0362 | MN337231 | MN336235 | MN366246 | MN509783 | Lambert et al. (2019) |
| *Hypoxydon fragiforme* | MUCL 51264<sup>T</sup> | KC477229 | KM186295 | KM186296 | KX271282 | Daranagama et al. (2015) |
| *Hypoxydon lienhwacheense* | MFLUCC 14-1231 | KU604558 | MK287550 | MK287563 | KU159522 | Sir et al. (2019) |
| *Pyrenopolyporus hunteri* | STMA07009 | – | – | KY624309 | – | Wendt et al. (2018) |
| *Idriella lunata* | CBS 204.56<sup>T</sup> | MH857584 | MH869129 | – | – | Vu et al. (2019) |
| *Jackrogersella multiformis* | CBS 119016<sup>T</sup> | KC477234 | KY610473 | KY624290 | KX271262 | Wendt et al. (2018) |
| *Kretzschmaria deusta* | CBS163.93 | – | – | KY624227 | – | Wendt et al. (2018) |
| *Kretzschmariella culmorum* | JDR88 | KX430043 | – | KX430045 | KX430046 | Fournier et al. (2018) |
| *Leptosillia cordyline* | MFLU 20-0056 | MT073349 | NG_068699 | MT023134 | MT023136 | Senanayake et al. (2020) |
| *Linosporeopsis ischnotheca* | LIF1 | MN818952 | MN818952 | MN820708 | MN820715 | Voglmayr and Beenken (2020) |
| *Linosporeopsis ochracea* | LIO3 | MN818958 | MN818958 | MN820714 | MN820721 | Voglmayr and Beenken (2020) |
| *Longiappendispora chromolaenae* | MFLU 20-0320<sup>T</sup> | MT214370 | MT214464 | – | – | Mapook et al. (2020) |
| *Lopadostoma americanum* | CBS 133211<sup>T</sup> | KC774568 | KC774568 | KC774525 | – | Jaklitsch et al. (2014) |
| *Lopadostoma dryophilum* | LG21 | NR_132028 | KC774570 | KC774526 | MF489023 | Jaklitsch et al. (2014) |
| *Lopadostoma fagi* | CBS 133206<sup>T</sup> | KC774575 | KC774575 | KC774531 | – | Jaklitsch et al. (2014) |
| *Lopadostoma gastrinum* | MFLU 17-0941 | MN337229 | MN336233 | MN366245 | – | Jaklitsch et al. (2014) |
| *Lopadostoma insulare* | CBS 133214<sup>T</sup> | KC774589 | KC774589 | KC774542 | – | Jaklitsch et al. (2014) |
| *Lopadostoma lechatii* | CBS 133694<sup>T</sup> | KC774590 | KC774590 | KC774543 | – | Jaklitsch et al. (2014) |
| *Lopadostoma quercicola* | CBS 133212<sup>T</sup> | KC774610 | KC774610 | KC774558 | – | Jaklitsch et al. (2014) |
| *Lopadostoma turgidum* | CBS 133207<sup>T</sup> | KC774618 | KC774618 | KC774563 | MF489024 | Jaklitsch et al. (2014) |
| *Lunatiannulus irregularis* | MFLUCC 14-0014<sup>T</sup> | KP297398 | KP340540 | KP340526 | KP406609 | Daranagama et al. (2015) |
| *Melanographium phoenicis* | MFLUCC 18-1481<sup>T</sup> | MN482677 | MN482678 | – | – | Hyde et al. (2020) |
| *Microdochium lycopodium* | CBS 125585<sup>T</sup> | JF440979 | JF440979 | KP859125 | KP859080 | Jaklitsch and Voglmayr (2012) |
| *Microdochium phragmitis* | CBS 285.71<sup>T</sup> | KP859013 | KP858949 | KP859122 | KP859077 | Jaklitsch and Voglmayr (2012) |
| *Monosporascus cannonballus* | MC1703 | JQ743053 | – | – | JQ804936 | Cavalcante et al. (2020) |
| *Muscodor thailandicus* | MFLUCC 17-2669<sup>T</sup> | MK762707 | MK762714 | MK791283 | MK776960 | Samarakoon et al. (2020) |
| *Muscodor ziziphi* | MFLUCC 17-2662 | MK762705 | MK762712 | MK791281 | MK776958 | Samarakoon et al. (2020) |
| *Nemania abortive* | BISH467<sup>T</sup> | GU292816 | – | GQ844768 | GQ470219 | Hsieh et al. (2010) |
| *Nemania beaumontii* | HAST405 | GU292819 | – | GQ844772 | GQ470222 | Hsieh et al. (2010) |
| *Nemania bipapillata* | HAST90080610 | GU292818 | – | GQ844771 | GQ470221 | Hsieh et al. (2010) |
| *Nemania diffusa* | HAST91020401 | GU292817 | – | GQ844769 | GQ470220 | Hsieh et al. (2010) |
| *Nemania illita* | YMJ236 | EF026122 | – | GQ844770 | EF025608 | Hsieh et al. (2010) |
| *Nemania macrocarpa* | WSP265<sup>T</sup> | GU292823 | MH874423 | GQ844776 | GQ470226 | Hsieh et al. (2010) |
| *Nemania maritima* | HAST89120401<sup>T</sup> | GU292822 | – | GQ844775 | GQ470225 | Hsieh et al. (2010) |
| *Nemania primolutea* | YMJ91102001<sup>T</sup> | EF026121 | – | GQ844767 | EF025607 | Hsieh et al. (2010) |
| *Nemania sp.* | BCC30850 | KX037428 | KX037429 | KX037430 | – | Kornsakulkarn et al. (2017) |
| *Nemania sphaerostoma* | JDR 261 | GU292821 | – | GQ844774 | GQ470224 | Hsieh et al. (2010) |
| *Neanthostomella fici* | MFLU 19-2765<sup>T</sup> | MW114390 | MW114445 | MW177711 | – | Tennakoon et al. (2021) |
| *Neanthostomella viticola* | MFLUCC 16-0243<sup>T</sup> | KX505957 | KX505958 | KX789496 | KX789495 | Daranagama et al. (2015) |
| *Neoeutypella baoshanensis* | MFLUCC 16-1002<sup>T</sup> | MT310662 | MT214618 | – | – | Phukhamsakda et al. (2020) |
| *Neoidriella desertorum* | CBS 985.72<sup>T</sup> | KP859048 | KP858985 | – | – | Hernandez-Restrepo et al. (2016) |
| *Obolarina persica* | YMJ 1461 | JX507807 | – | JX507792 | JX507796 | Mirabolfathy et al. (2013) |
| *Paraeutypella citricola* | HUEFS 194248 | KM396616 | KM407630 | – | KR534602 | de Almeida et al. (2016) |
| *Paraeutypella vitis* | UCD2428TX | FJ790851 | – | – | GU294726 | Urbez-Torres et al. (2009) |
| *Paraxylaria rosacearum* | TASM 6132<sup>T</sup> | MG828941 | MG829050 | – | – | Wanasinghe et al. (2018) |
| *Pedumispora rhizophorae* | BCC44877 | KJ888853 | KJ888850 | – | – | Klaysuban et al. (2014) |
| *Peroneutypa diminutiasca* | MFLUCC 17-2144<sup>T</sup> | MG873479 | MG873473 | – | MH316765 | Shang et al. (2018) |
| *Peroneutypa diminutispora* | HUEFS 192196<sup>T</sup> | KM396647 | KM407664 | – | – | de Almeida et al. (2016) |
| *Peroneutypa mackenziei* | MFLUCC 16-0072<sup>T</sup> | KY283083 | KY283079 | – | KY706363 | Shang et al. (2017) |
| *Peroneutypa mangrovei* | PUFD526<sup>T</sup> | MG844286 | MG844278 | – | MH094409 | Phookamsak et al. (2019) |
| *Peroneutypa scoparia* | MFLUCC 11-0615 | KU940152 | KU863140 | KU940180 | – | Dai et al. (2016) |
| *Pirozynskiomyces sinensis* | 2184417 | KY994106 | KY994107 | – | – | Li et al. (2017) |
| *Podosordaria mexicana* | WSP176 | GU324762 | – | GQ853039 | GQ844840 | Hsieh et al. (2010) |
| *Poronia punctate* | CBS 656.78<sup>T</sup> | KT281904 | KY610496 | KY624278 | KX271281 | Senanayake et al. (2015) |
| *Pseudoanthostomella conorum* | CBS 119333 | EU552099 | EU552099 | – | – | Marincowitz et al. (2008) |
| *Pseudoanthostomella delitescens* | MFLUCC 16-0477 | KX533451 | KX533452 | KX789491 | KX789490 | Daranagama et al. (2015) |
| *Pseudoanthostomella pini-nigrae* | MFLUCC 16-0478<sup>T</sup> | KX533453 | KX533454 | KX789492 | – | Daranagama et al. (2015) |
| *Pseudoceratocladium polysetosum* | FMR10750 | KY853430 | KY853490 | – | – | Hernandez-Restrepo et al. (2017) |
| *Pseudocircinotrichum papakurae* | CBS101373 | KR611876 | KR611897 | – | – | Crous et al. (2015) |
| *Pyrenomyxa morgani* | CBS 118186<sup>T</sup> | MH863028 | MH874581 | – | – | Vu et al. (2019) |
| *Pyrenopolyporus hunteri* | MUCL 52673<sup>T</sup> | KY610421 | KY610472 | KY624309 | KU159530 | Wendt et al. (2018) |
| *Pyriiformiascoma trilobatum* | MFLUCC 14-0012<sup>T</sup> | KP297402 | KP340543 | KP340530 | KP406613 | Daranagama et al. (2015) |
| *Requienella fraxini* | CBS 140475<sup>T</sup> | NR_138415 | KT949910 | – | – | Jaklitsch et al. (2016) |
| *Requienella seminuda* | CBS 140502<sup>T</sup> | NR_154630 | KT949912 | MK523300 | – | Jaklitsch et al. (2016) |
| *Rhopalostroma angolense* | CBS 126414 | FN821965 | KM186298 | KM186297 | KM186299 | Stadler et al. (2010a) |
| *Rosellinia aquila* | MUCL 51703 | KY610392 | KY610460 | KY624285 | KX271253 | Wendt et al. (2018) |
| *Rosellinia Chiangmaiensis* | MFLUCC 15-0015<sup>T</sup> | KU246226 | KU246227 | – | – | Daranagama et al. (2015) |
| *Rosellinia corticium* | MUCL 51693 | KY610393 | KY610461 | KY624229 | KX271254 | Wendt et al. (2018) |
| *Rosellinia limonispora* | CBS 382.86 | KR611876 | KR611897 | – | – | Hartley (2020) |
| *Rosellinia merrillii* | HAST89112601 | GU300071 | – | GQ844781 | GQ470229 | Hsieh et al. (2010) |
| *Rosellinia thelena* | CBS 400.61 | KF719202 | KF719215 | – | – | Wu et al. (2025) |
| *Ruwenzoria pseudoannulata* | MUCL 51394<sup>T</sup> | KY610406 | KY610494 | KY624286 | KX271278 | Wendt et al. (2018) |
| *Sarcoxyloa compunctum* | CBS 359.61 | KT281903 | KT281898 | KY624230 | KX271255 | Senanayake et al. (2015) |
| *Selenodriella cubensis* | CBS 683.96<sup>T</sup> | KP859053 | KP858990 | – | – | Hernandez-Restrepo et al. (2017) |
| *Seynesia erumpens* | SMH 1291 | – | AF279410 | AY641073 | – | Bhattacharya et al. (2020) |
| *Sporidesmina malabarica* | NN43118 | – | DQ408580 | – | – | Bhattacharya et al. (2020) |
| *Sporidesmina polypora* | NN47796 | OL627678 | DQ408569 | DQ435087 | – | Delgado et al. (2025) |
| *Stilbohypoxyton elaeidicola* | HAST 94082615 | GU322440 | – | GQ844827 | GQ495933 | Hsieh et al. (2010) |
| *Stilbohypoxyton elaeidis* | MFLUCC 15-0295b | MT496746 | MT496756 | – | – | Konta et al. (2020) |
| *Stromatoneurospora phoenix* | BCC82040 | MT703666 | MT735133 | MT742605 | MT700438 | Becker et al. (2020) |
| *Thamnomycetes dendroideus* | CBS 123578<sup>T</sup> | FN428831 | KY610467 | KY624232 | KY624313 | Stadler et al. (2010b) |
| *Theissenia cinerea* | YMJ 90071615<sup>T</sup> | EF026128 | – | JX507793 | EF025613 | Hsieh et al. (2010) |
| *Thuemenella cubispora* | CBS 119807 | JX658531 | EF562508 | – | – | Stadler et al. (2014) |
| *Vamsapriya bambusicola* | MFLUCC 11-0477<sup>T</sup> | KM462835 | KM462836 | KM462834 | KM462833 | Dai et al. (2014) |
| *Vamsapriya breviconidiophora* | MFLUCC 14-0436<sup>T</sup> | MF621584 | MF621588 | – | – | Li et al. (2017) |
| *Vamsapriya indica* | MFLUCC 12-0544 | KM462839 | KM462840 | KM462841 | KM462838 | Dai et al. (2014) |
| *Vamsapriya khunkonensis* | MFLUCC 11-0475<sup>T</sup> | MW240620 | MW240549 | KM462829 | KM462828 | Dai et al. (2014) |
| *Vesiculozygospodium echinosporum* | CPC 35607<sup>T</sup> | MT223869 | MT223940 | – | – | Crous et al. (2020) |
| *Virgaria nigra* | CBS 128006 | MH864744 | MH876180 | – | – | Vu et al. (2019) |
| *Xenoanthostomella chromolaenae* | MFLUCC 17-1484<sup>T</sup> | MN638863 | MN638848 | MN648729 | – | Hyde et al. (2020) |
| *Xenoanthostomella cycadis* | CPC17285 | KJ869121 | KJ869178 | – | – | Crous et al. (2014) |
| *Xylaria allantoidea* | BCC22746 | MT703671 | MT735141 | MT742610 | – | Becker et al. (2020) |
| *Xylaria atosphaerica* | HAST91111214 | GU322459 | – | GQ848342 | GQ495953 | Hsieh et al. (2010) |
| *Xylaria digitate* | HAST919 | GU322456 | – | GQ848338 | GQ495949 | Hsieh et al. (2010) |
| *Xylaria heliscus* | HAST88113010 | GU324742 | – | GQ848355 | GQ502691 | Hsieh et al. (2010) |
| *Xylaria hypoxylon* | CBS122620<sup>T</sup> | AM993141 | KM186301 | KM186302 | KM186300 | Persoh et al. (2009) |
| *Xylaria longipes* | CBS 148.73 | MH860649 | MH872351 | KU684280 | KU684204 | Vu et al. (2019) |
| *Xylaria polymorpha* | MUCL 49884 | KY610408 | KY610464 | KY624288 | KX271280 | Wendt et al. (2018) |
| *Xylaria regalis* | HAST92072001 | GU324744 | – | GQ848357 | GQ502693 | Hsieh et al. (2010) |
| *Zygospodium minus* | HKAS 99625<sup>T</sup> | MF621586 | MF621590 | – | – | Li et al. (2017) |
| *Zygospodium oscheoides* | MFLUCC 14-0402<sup>T</sup> | MF621585 | MF621589 | – | – | Li et al. (2017) |

### Phylogenetic analyses

Newly generated sequences were subjected to BLASTn searches against the GenBank database (http://www.ncbi.nlm.nih.gov/) to determine preliminary taxonomic affiliations. Additional sequences were obtained from GenBank based on BLAST results and previous studies. A concatenated dataset of LSU, ITS, *RPB2* and *TUB2* sequences was assembled, comprising 202 strains. *Leptosillia cordylines* (MFLU 20-0056) and *Delonicicola siamense* (MFLUCC 15-0670) were selected as the outgroup.

Individual gene alignments were generated using the online version of MAFFT v. 7.0362 (Katoh et al. 2019) with default parameters and manually refined in BioEdit 7.1.3 (Hall 1999) where necessary. Maximum Likelihood (ML) analysis was performed with RAxML (Stamatakis and Alachiotis 2010) implemented in raxmlGUIv.1.5 (Silvestro and Michalak 2012) using the ML + rapid bootstrap setting and the GTR + I + G model of nucleotide substitution with 1,000 rapid bootstrap replicates. For Bayesian inference (BI), the optimal substitution model (GTR+G+I) was determined using MrModeltest software v. 2.2 (Nylander 2004). BI analysis was conducted in MrBayes v. 3.2.6 (Ronquist et al. 2012) with four simultaneous Markov chain Monte Carlo chains from random trees over 3M generations (standard deviation of split frequencies < 0.01), sampling every 500^th^ generations. The distribution of log-likelihood scores was detected to check whether sampling was in the stationary phase, and Tracer v1.5 was used to check if further runs were required to reach convergence (Rambaut and Drummond 2007). The first 20% of sampled trees were discarded as burn-in before calculating a consensus tree and posterior probabilities. Resultant phylogenetic trees were visualized using FigTree v.1.4 (Rambaut 2012).

### De novo genome assembly and Genome sequencing of *Dianjunus rex*

Whole-genome and transcriptome sequencing were performed on three stromata of *Dianjunus rex* collected from the wild. Internal tissue was excised from each stromata. High-quality genomic DNA was extracted from each sample using the CTAB method (Porebski et al. 1997). Total RNA was extracted with TRIzol® Reagent (Invitrogen) extraction kit, followed by purification using the Plant RNA Purification Reagent (Invitrogen) purification kit. Short DNA and RNA reads were generated for each sample on the Illumina HiSeq4000 platform (San Diego, CA, USA). These reads were then trimmed and filtered using fastp (v0.23.2) (Chen et al. 2018) with parameters ‘-f 10 and -F 10’ to remove adapter sequences and low-quality reads. For paired-end DNA sequencing data, after converting the clean data into a FASTA file, 10,000 reads were randomly selected from each file and aligned against the NCBI Nucleotide (NT) database, indicating no significant contamination. Finally, in total, we obtained 49 Gb of clean data for DNA and 51 Gb for RNA, respectively. For genome assembly, the filtered DNA reads from each sample were assembled independently using by SPAdes (v4.2.0) (Bankevich et al. 2012). Genome completeness was assessed by Benchmarking Universal Single-Copy Orthologs (BUSCO, v5.2.2) (Simao et al. 2015) with the ascomycota_odb10 orthologous gene set.

### Annotation of repetitive sequences and protein-coding genes

RepeatModeler (v2.0.2) (Flynn et al. 2020) was used to build the *de novo* repeat libraries and then sequence for the assembly were aligned to the *de novo* repeat libraries and known repeat databases to identify and mask repetitive elements, using RepeatMasker (v4.1.2) (Chen 2004). Protein-coding genes in each genome were predicted using GETA (v2.6.1) pipeline (https://github.com/chenlianfu/geta), integrating evidence from homology-based, ab initio, and RNA-seq-based prediction approach. For homology-based prediction, protein-coding sequences of *Biscogniauxia marginata, Annulohypoxylon maeteangense*, *Daldinia caldariorum*, *Hypoxylon argillaceum*, *Astrocystis sublimbata*, *Durotheca rogersii*, *Kretzschmaria deusta*, *Nemania abortiva*, *Poronia punctata*, and *Xylaria curta* were download from the Joint Genome Institute (JGI) (Franco et al. 2022) and were used for gene prediction using GeneWise (v2.4.1) (Birney et al. 2004). The ab initio prediction was conducted using Augustus (v3.4.0) (Stanke et al. 2006). RNA-seq reads were aligned to the genome assembly using HISAT2 (v2.1.0) (Kim et al. 2015), and protein-coding regions were predicted using TransDecoder (v5.5.0) (https://github.com/TransDecoder/TransDecoder). The predictions from the three methods were then integrated and filtered against the PFAM database.

### Phylogenetic tree construction and divergence time estimation

Protein sequences of 63 species in *Pezizomycotina* were downloaded from JGI Mycocosm database, combined with three samples generated in this study, resulting in 66 samples for phylogenetic analysis. Single-copy orthologous proteins conserved across all 66 samples were retrieved from the results of BUSCO assessment. A total of 302 single copy orthologs were then aligned using MAFFT (v7.407) (Katoh et al. 2002) with default parameters, and the alignments were concatenated sequentially to construct a super matrix. The phylogenetic tree was constructed from this super matrix using IQ-TREE3 (v3.0.1) (Wong et al. 2025), with *Aspergillus niger* and *A. tetrazonus* serving as the outgroup. Branch supports were assessed using bootstrap approach with 1,000 replicates. The species tree was visualized using the web application tvBOT (Xie et al. 2023).

To further cross-verify the topological stability across different ortholog selection workflows, an alternative dataset was constructed. Orthologous groups across the same 66 samples were clustered using OrthoFinder (v3.1.3) with the DIAMOND alignment mode (-S diamond) and MAFFT for sequence alignment (-M msa -A mafft), yielding 345 single-copy orthologous gene sets. The amino acid sequences of each orthogroup were aligned using MAFFT (v7.407) with automatic parameter selection (--auto). To eliminate poorly aligned or highly divergent regions, the alignments were systematically trimmed using trimAl (v1.4.rev15) with a gap threshold of 0.3 (-gt 0.3), ensuring that sites with more than 70% gaps were removed. Individual maximum-likelihood gene trees were then inferred from the trimmed alignments using IQ-TREE3 (v3.0.1) under the auto-selected best-fit evolutionary models (MFP) with 1,000 ultrafast bootstrap replicates, utilizing the same outgroup (*Aspergillus niger* and *A. tetrazonus*). Finally, these 345 gene trees were used as input for ASTRAL (v5.7.1) to reconstruct the alternative coalescent-based species tree.

The MCMCtree module of PAML package (v4.9i) (Yang et al. 2007) was used to estimate each speciation times for these 66 *Pezizomycotina* samples. The phylogenetic tree constructed by IQ-TREE3 served as a guide tree, with the divergence times from TimeTree (Kumar et al. 2017) used as calibration points: *Aspergillus tetrazonus* and *Pyricularia oryzae* (233.8∼367.0 MYA), *Pseudomassariella vexata* and *Microdochium trichocladiopsis* (143.3∼164.1 MYA), *Microdochium bolleyi* and *Hypoxylon fuscum* (103.4∼318.2 MYA), and *Colletorichum cereale* and *Verticillium longisporum* (85.0∼139.4 MYA).

### Identification and Analysis of Carbohydrate-Active Enzymes

CAZymes profiles were identified in the proteomes of all the 66 *Pezizomycotina* samples. BLAST and hidden Markov model (HMM) searches were performed simultaneously using the proteomes as queries to detect putative CAZymes. For the BLAST searches, protein sequences of queries were used as inputs in the BLASTp tool against dbCAN database (v11) (Yin et al. 2012). The resulting hits were filtered by e-value (1e-5) and identity threshold (20%). For the HMM search, Hmmscan program in the HMMER package was used to match the family-specific HMM profiles of CAZymes within dbCAN database (v11). The candidate genes obtained from both the BLAST and HMM searches were merged and only the overlapping hits were retained. The raw sequencing data have been deposited in the National Center for Biotechnology Information (NCBI) under the BioProject accession number of PRJNA1265937. All genome assemblies and gene annotation results have been archived in the figshare database and can be accessed at https://doi.org/10.6084/m9.figshare.29153135.v2. The multilocus alignment of this study can be accessed via https://doi.org/10.6084/m9.figshare.32611812.

## Results

### Phylogeny

All individual trees generated under different criteria and from single gene datasets were essentially similar in topology and not significantly different from the tree generated from the concatenated dataset (not discussed herein). Maximum likelihood analysis of new strains generated in this study and other related taxa in this study with 1,000 bootstrap replicates yielded the best ML tree (Fig. 1) with the likelihood value of ln: –6728.286175 and the following model parameters: alpha: 0.191444; Π(A): 0.244188, Π(C): 0.230054, Π(G): 0.301322 and Π(T): 0.224436. The substitution rates are as: AC: 0.763664, AG: 2.779076, AT: 1.349374, CG: 0.639806, CT: 6.128235, GT: 1.0. The 10,000 trees resulted in Bayesian analysis after 5,000,000 generations and the first 20% trees were discarded. The remaining trees were used for calculating posterior probabilities in the majority rule consensus tree. Maximum likelihood bootstrap values ≥50% and Bayesian inference (BI) ≥0.9 are given at each node. Tree topologies of the ML and Bayesian analyses were similar to each other and there are no any significant differences. Both topologies showed well-supported, 15 major clades representing families in *Xylariales*. These new fungal collections grouped as a sister clade to *Graphostromataceae* with ML/BI= 100%/1.00 support.

**Fig. 1.**
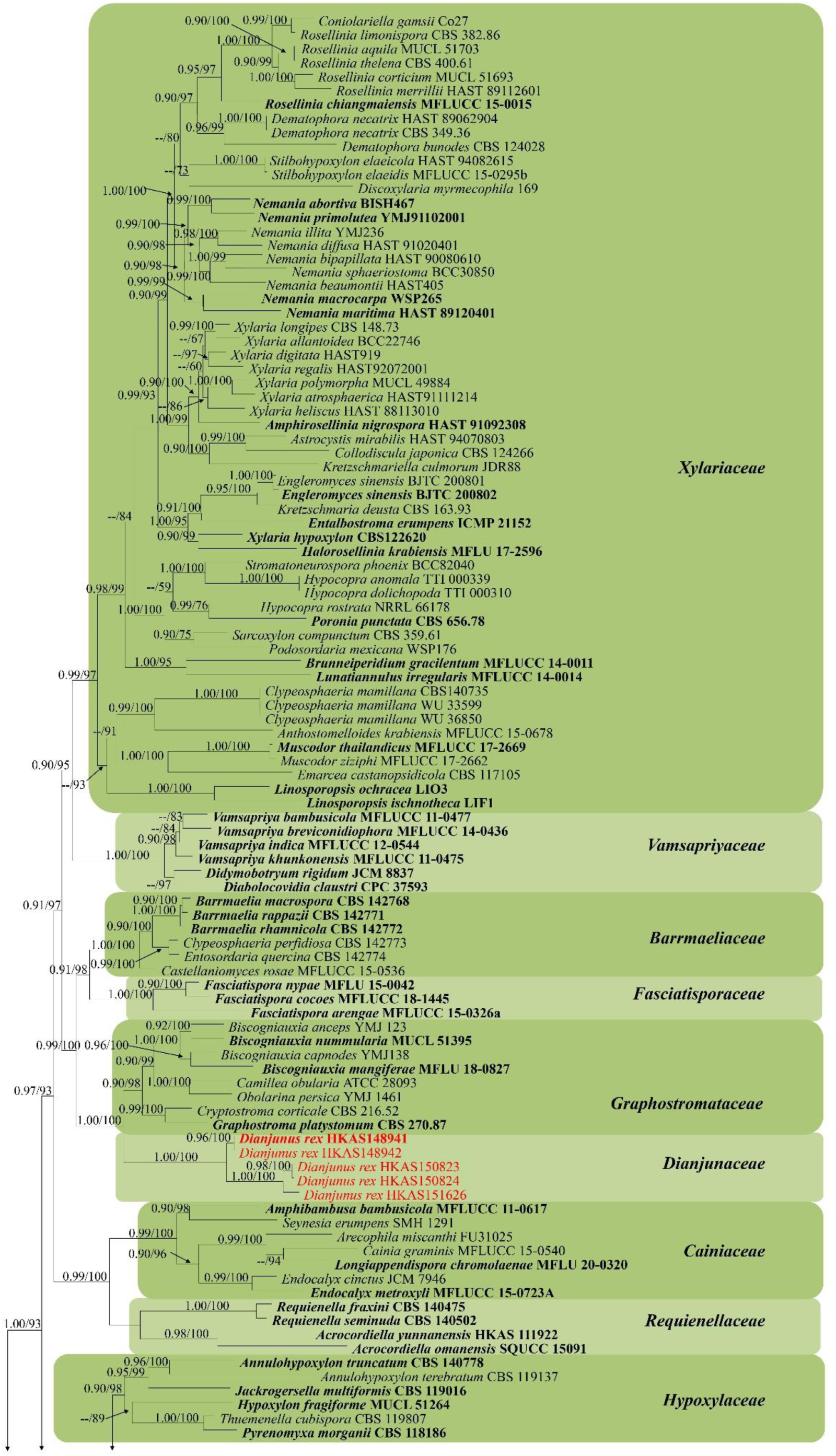
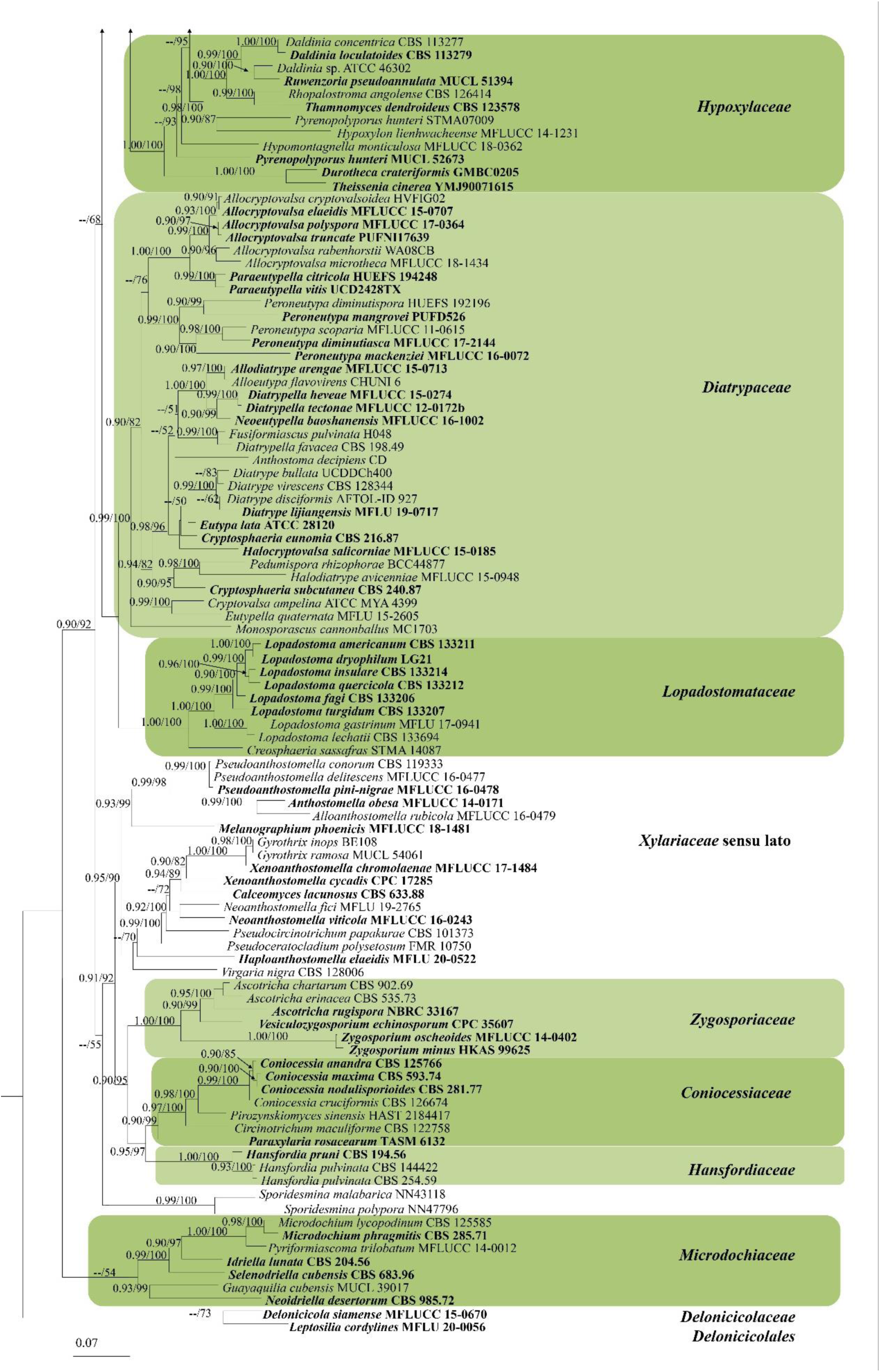
Phylogenetic tree inferred from maximum likelihood analysis of the combined ITS, LSU, *TUB2*, and *RPB2* sequence alignment. Nodes are labelled with maximum likelihood bootstrap support values ≥50%, and Bayesian posterior probabilities ≥0.90. The tree was rooted with *Leptosillia cordylines* (MFLU 20-0056) and *Delonicicola siamense* (MFLUCC 15-0670). Ex-type cultures are denoted in bold, and sequences newly generated in this study are highlighted in red.

### Phylogenomic tree construction and divergence time estimation

The genome assembly sizes of sample HKAS148941, HKAS150823 and HKAS151626 are 31.09 Mb, 32.01 Mb, and 30.48 Mb with a contig N50 of 47.63 Kb, 42.61 Kb and 76.62 Kb respectively (Table 3). Genome completeness was revealed that at least 96.19%, 95.96% and 96.19% single-copy BUSCO genes were identified in the assembly (Fig. 2, Table 3).

**Table 3.** Genome assembly statistics of *Dianjunus rex*.

| Sample | Contig N50 (Kb) | Contig Num | Total Length (Mb) | Protein Coding Genes |
| --- | --- | --- | --- | --- |
| HKAS148941 | 47.63 | 1471 | 31.19 | 7793 |
| HKAS150823 | 42.61 | 1437 | 32.01 | 6753 |
| HKAS151626 | 76.62 | 889 | 30.48 | 6779 |

**Fig. 2.**
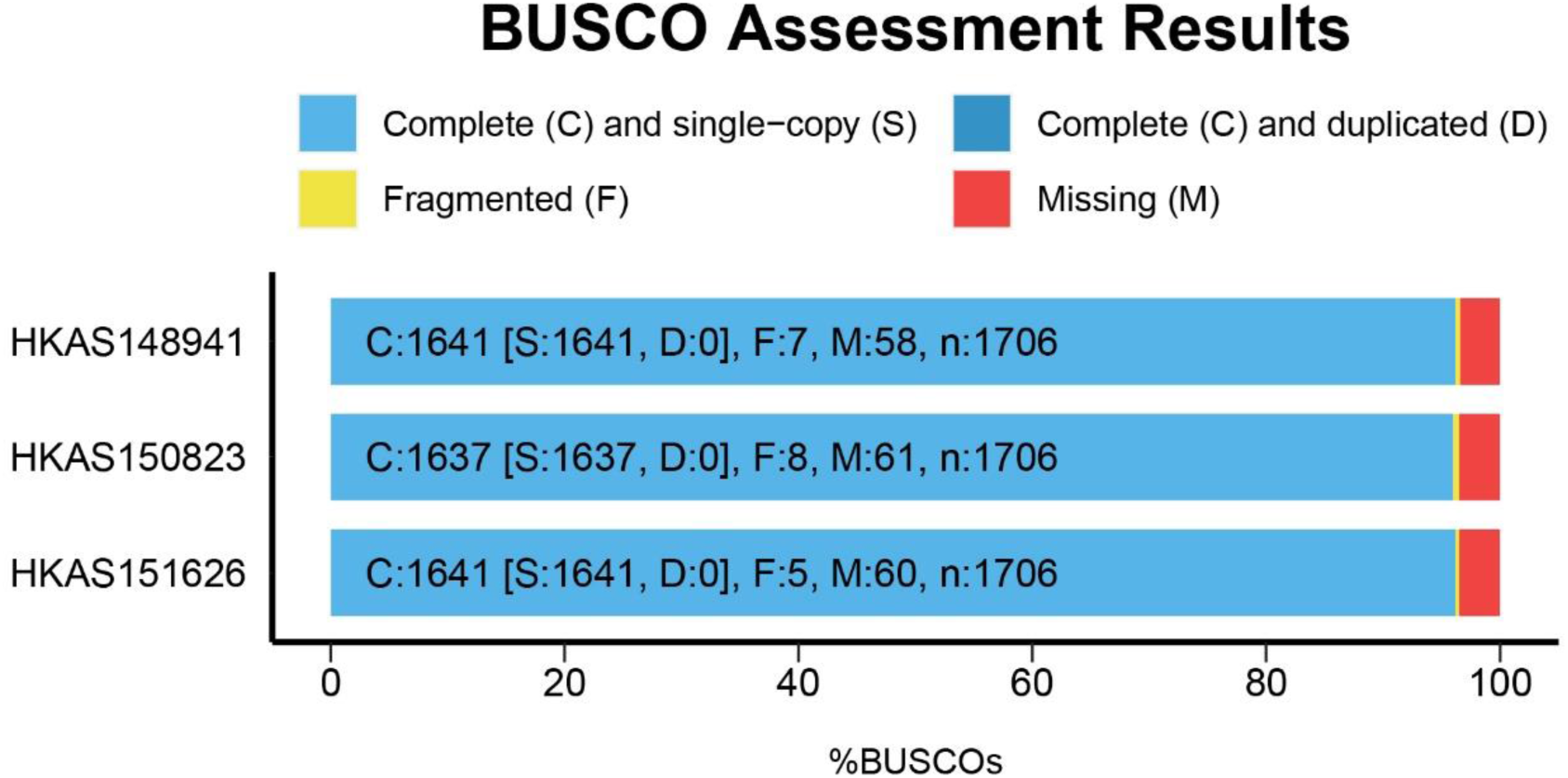
BUSCO assessments of the genome assembly and proteins of *Dianjunus rex*.

**Fig. 3.**
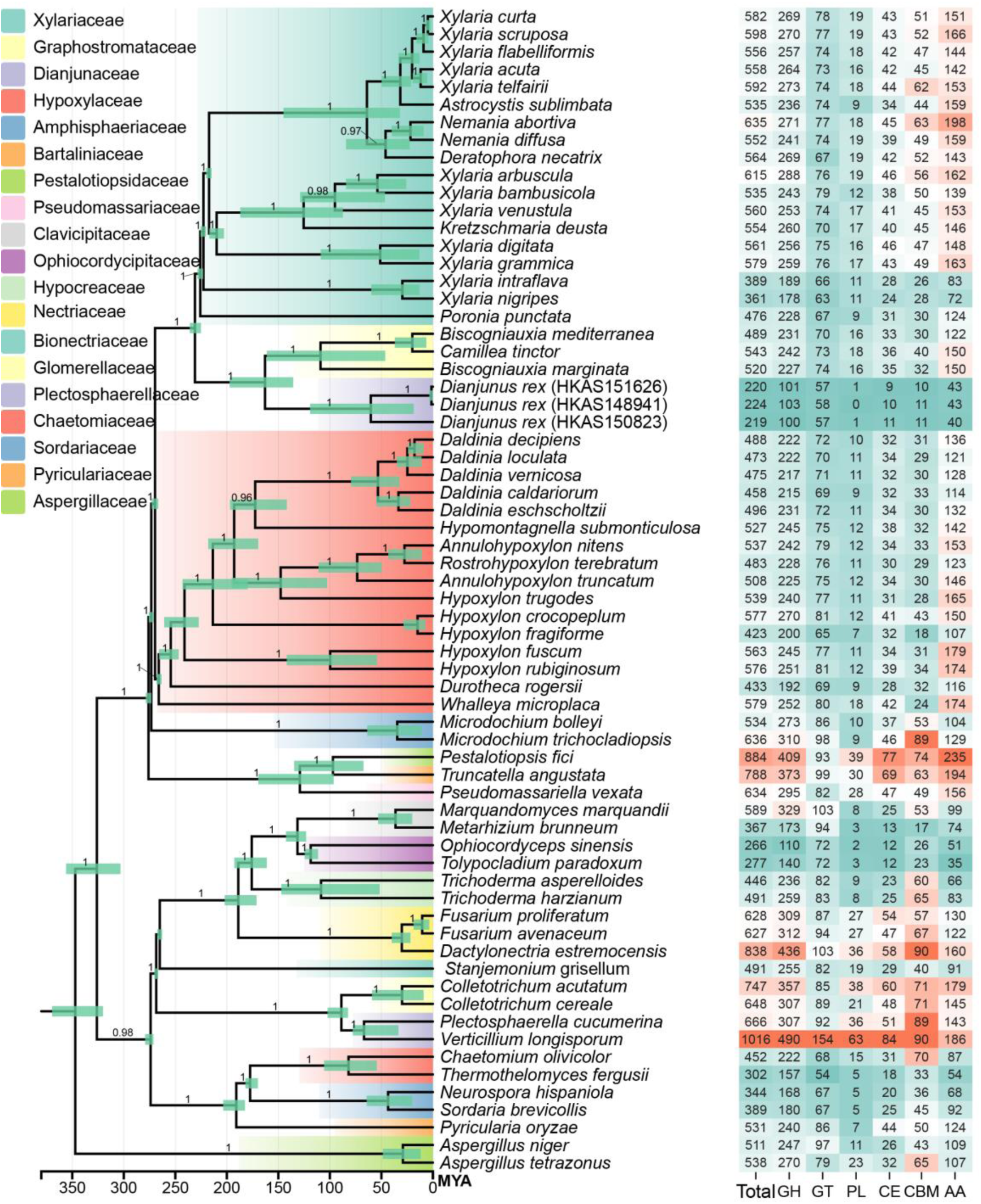
Phylogenomic tree inferred from a genome-scale dataset comprising 66 *Pezizomycotina* species with *Aspergillus niger* and *A. tetrazonus* as outgroup taxa. Devonian (420–360 Ma), Carboniferous (360–300 Ma), Permian (300–250 Ma), Triassic (250–200 Ma), Jurassic (200–145 Ma), Cretaceous (145–65 Ma), Paleogene (65–23 Ma), Neogene (23–2.6 Ma).

### Taxonomy

***Dianjunaceae* F.Q. Yu, Pérez-Moreno & Senan., fam. nov**

Index Fungorum number: IF904757

### Etymology

Based on the type genus *Dianjunus*.

### Type genus

***Dianjunus*** F.Q. Yu, Pérez-Moreno & Senan.

### Description

Teleomorph: Stromata upright, fleshy, variable in size, shape ranging from cylindrical, fusiform, turbinate to palm-shaped, terete to flattened, simple to branched from the base, sometime arising from long rooting stipes, with flattened sterile apices. Outer crust coriaceous, immersed part in soil black, aerial part fertile, surface covered with a long persistent, hairy, whitish yellow fibrous substance, whitish in young stromata, turning progressively dark yellow at maturity, ostiole openings appear as black dots. Immersed part sterile consisted with stipes, cylindrical, black, smooth. Internal tissues solid, homogenous to fibrous, spongy, off-white to yellow, with a slightly darker core in aged specimens. Perithecia aggregated, immersed in stromata, subglobose, coriaceous, papillate, ostiolate. Ostioles raised-discoid, black, with a cylindrical papilla at center. Paraphyses sparse, hypha-like, narrowing towards the ends, hyaline, septate, slightly constricted at the septa, unbranched. Asci (6–)8-spored, unitunicate, cylindrical, long-stipitate, total length, with apical apparatus, appearing bilobed, bluing in Melzer’s reagent, reddish brown in Lugol’s solution, turning blue in Lugol’s solution after treat with 3% KOH. Ascospores overlapping uniseriate in the ascus, globose to oval or fusiform with acute ends, hyaline when young, becoming brown at maturity, with single, central guttule or with a large central and two, small polar guttules, smooth, thick-walled, with a conspicuous, S-shaped germ slit one-half to four-fifths spore length on the flattened side. Anamorph: nodulisporium-like anamorph with periconiella-like branching patterns.

### Notes

In the multilocus phylogeny of ITS, LSU, *TUB2*, and *RPB2* sequences showed that our new fungal collections clustered together forming a well-supported, distinct sister clade to *Graphostromataceae* in *Xylariales*. However, morphological characteristics of our collections are significantly different from species in this family. Further, only few genera in *Graphostromataceae* produce upright stromata such as *Camillea*, and *Biscogniauxia*. The phylogenomic analyses also confirmed the placement of these collections sister to *Graphostromataceae* in *Xylariales*. Therefore, we introduce this unidentified clade as a new family *Dianjunaceae*.

***Dianjunus*** F.Q. Yu, Pérez-Moreno & Senan., gen. nov

**Index Fungorum number**: IF904733

### Etymology

The generic name derived from the “Dian Kingdom”, a historical society situated in the region of present-day Yunnan Province in south-western China where the samples were collected. The epithet is combined with the Chinese word “jūn”, meaning ’mushroom’ or ’fungus’.

### Type species

*Dianjunus rex* F.Q. Yu, Pérez-Moreno & Senan.

### Description

Arise from roots of dying host plants. Teleomorph: Stromata upright, fleshy, variable size and shape, ranging from cylindrical, fusiform, turbinate to palm-shaped, terete to flattened, simple to branched from the base, or at the top, arising from long rooting stipes, with flattened sterile apices. Outer crust coriaceous, black, ostiole openings appear as black dots. Aerial part fertile, surface covered with a long persistent, powdery, whitish yellow fibrous substance, whitish in young stromata, turning progressively dark yellow at maturity. Immersed part sterile consisted with stipes, cylindrical, black, no powdery substance, smooth. Interior tissues solid, homogenous to fibrous, spongy, off-white to yellow, with a slightly darker core in aged specimens, discolouration absence with 5% KOH. Ascomata perithecial, aggregated, scattered, immersed in stromata, subglobose, coriaceous, papillate, ostiolate. Ostioles raised-discoid, black, with a central, cylindrical papilla. Peridium dark brown, turning subhyaline inwardly, contain cells of *textura porrecta* mixed with *textura angularis*, evenly ticked. Paraphyses sparse, hypha-like, narrowing towards the ends, hyaline, septate, slightly constricted at the septa, unbranched, longer than asci, embedded in mucilaginous matrix. Asci (6–)8-spored, unitunicate, cylindrical, long-stipitate, total length, with apical apparatus, appearing bilobed, amyloid, cuboid in Melzer’s reagent, reddish brown in Lugol’s solution, turning blue in Lugol’s solution after treat with 3% KOH, inverted trapezoid. Ascospores overlapping uniseriate in the ascus, globose to oval or fusiform with acute ends, hyaline when young, becoming brown at maturity, with single, central guttule or with a large central and two, small polar guttules, smooth, thick-walled, with a conspicuous, S-shaped germ slit one-half to four-fifths spore length on the flattened side. Anamorph: nodulisporium-like with periconiella-like branching patterns.

### Notes

In the phylogenetic analyses of combined ITS, LSU, *TUB2*, and *RPB2* locus and phylogenomic analyses shown that these five collections (HKAS150824, HKAS151626, HKAS150823, HKAS148941, HKAS148942) clustered forming a well-supported clade (Fig. 1). Morphological characters of collections in this clade are significantly different from sistering clades. Therefore, we establish this clade as *Dianjunus gen. nov*.

***Dianjunus rex* F.Q. Yu, Pérez-Moreno & Senan., sp. nov.**

Index Fungorum number: IF904732

Figs. 4–6

**Fig. 4.**
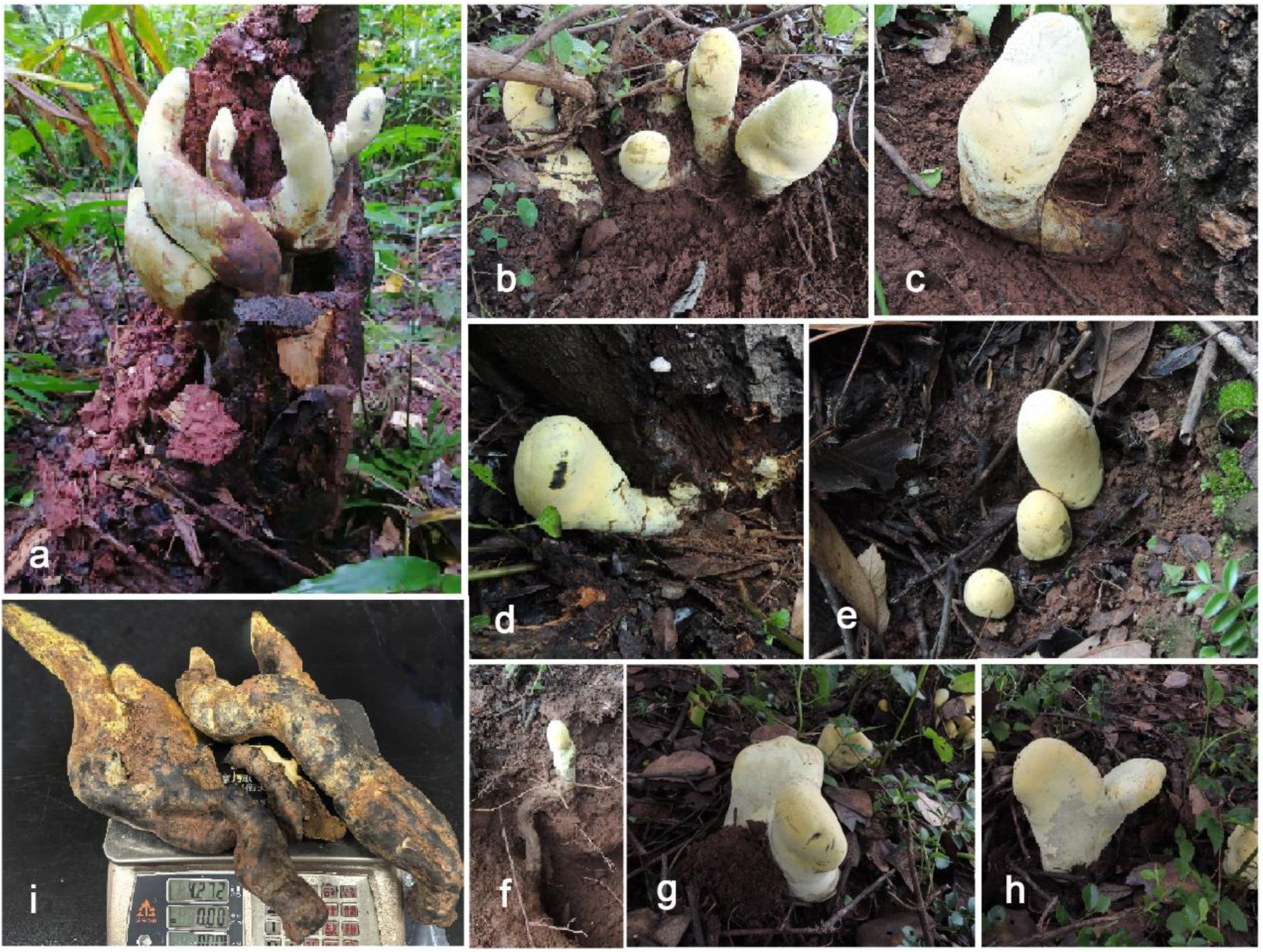
Stromata of *Dianjunus rex* at the field. **a, c, d, f** Stromata attached to host roots. **b, e, g, h** Stromata around host plant. **a** Collectors. **i** Total weight of largest bilobed-stromata.

**Fig. 5.**
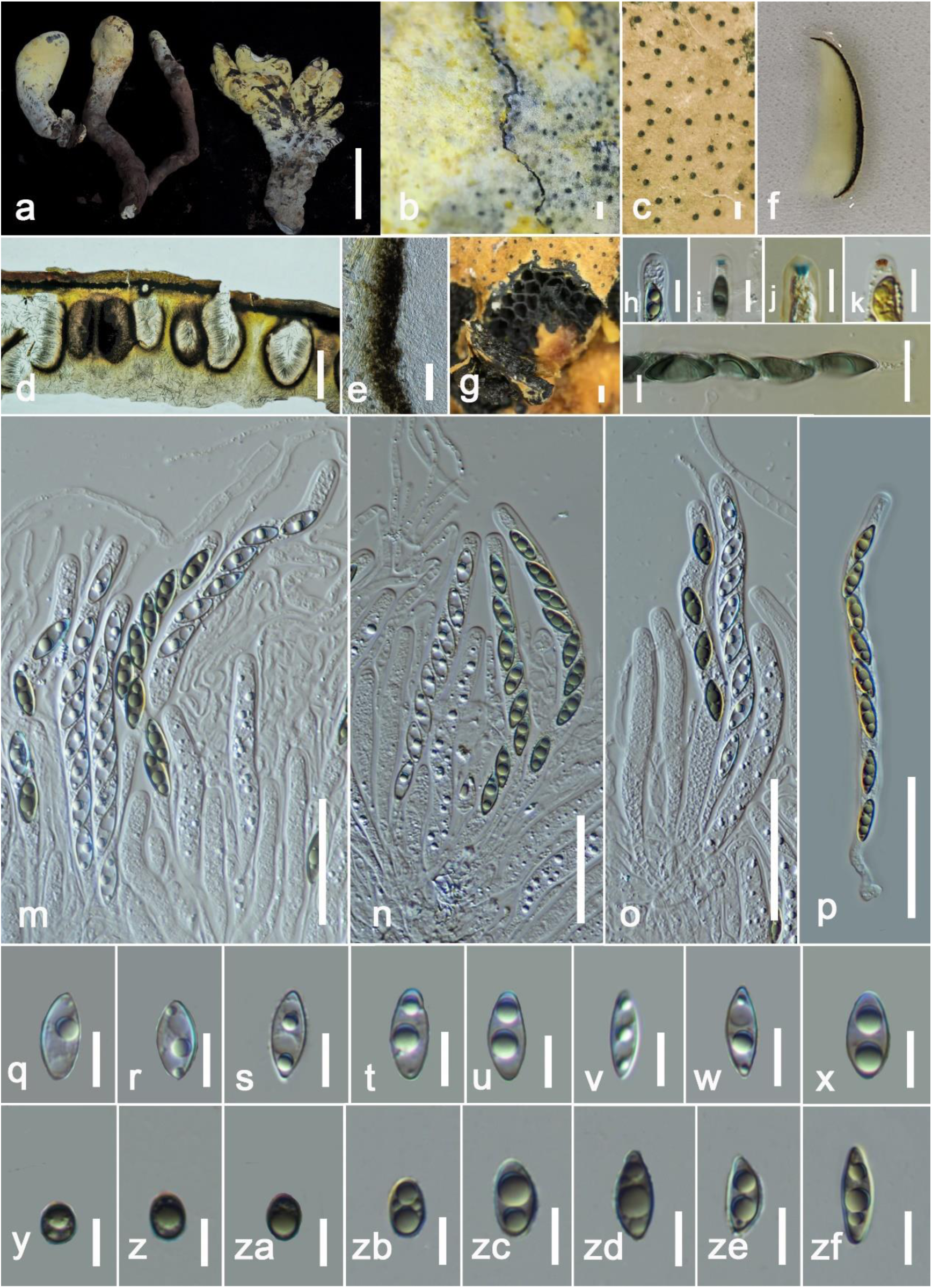
*Dianjunus rex* (HKAS148941, holotype). **a** Stromata. **b** Yellow powdery layer on outer crust covering ostioles. **c** Ostiole opening on stromatic surface. **d** Vertical cross section of ascoma. **e** Peridium. **f** Stromatic tissues in 5% KOH. **g** Horizontal section through the stromata. **h** Apical ring in water. **i** Apical ring in Melzer’s regent. **j** Apical ring in Lugol’s regent after pre-treat with 3% KOH. **k** Apical ring in Lugol’s regent. **l** S-shaped germ-slits in ascospores in Melzer’s regent. **m-p** Asci in water. **q-zf** Ascospores in water. Scale bars: a = 15 cm, b, c, e = 100 µm, d = 750 µm, g = 250 µm, l = 20 µm, m-p = 50 µm, h-k, q-zf = 10 µm.

**Fig. 6.**
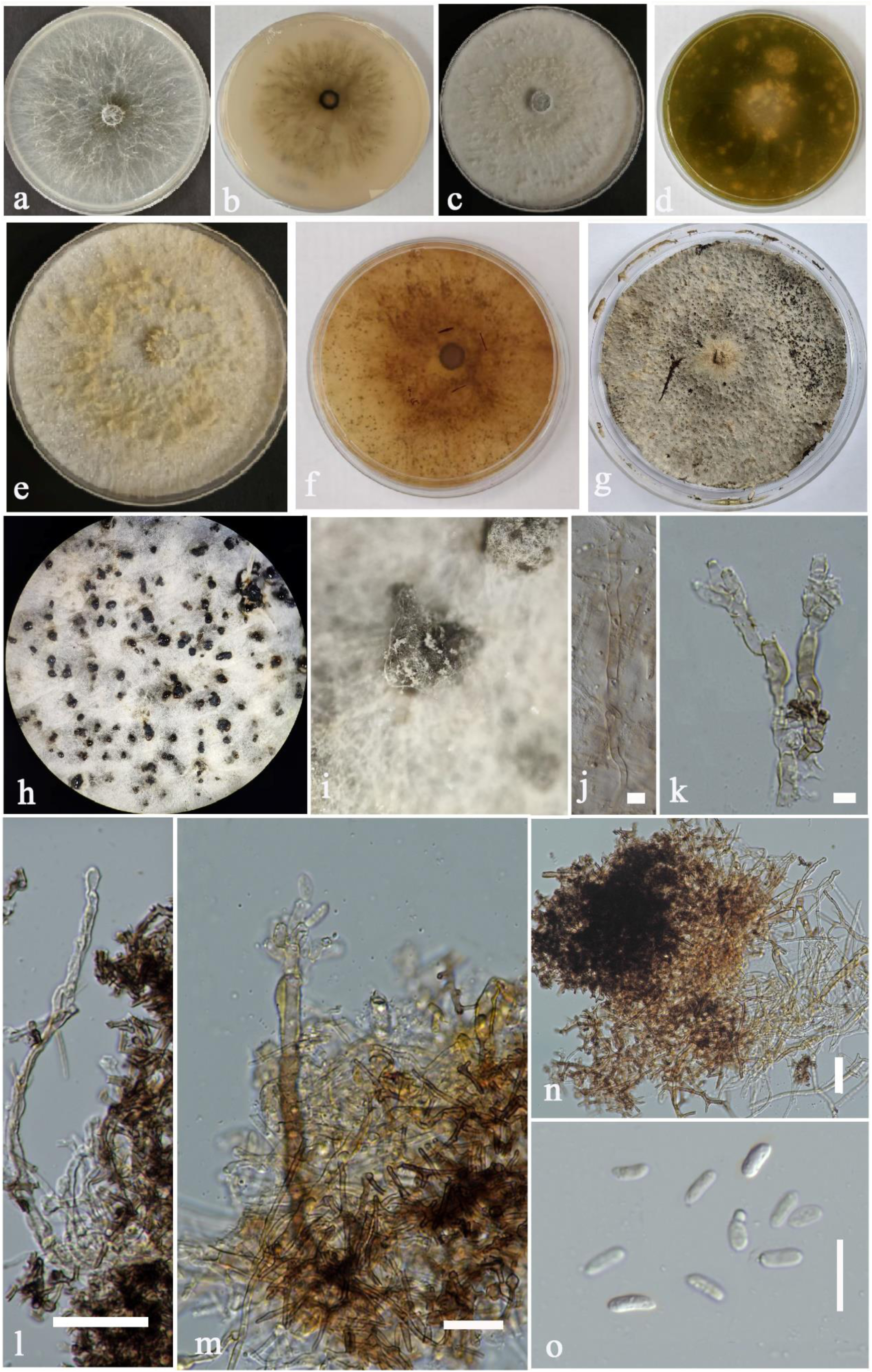
Cultures and anamorph of *Dianjunus rex*. (GBWSCC202508860). **a, c, e** Surface view of culture on MEA, OMA, PDA respectively after a week. **b,d,f** Reverse view of culture on MEA, OMA, PDA after a week. **g, h** 90 days old, sporulated culture on PDA. **i** Close-up view of spore-baring structures. **j** Mycelia. **k-m** Conidiophores, Conidiogenous cells attached with conidia. **n** Specialized mycelial cells which conidiophores are derived. **o** Conidia. Scale bars: l-n = 20 µm, j,k,o = 10 µm.

### Etymology

The species epithet *rex* (Latin = king), referring to the large size and majestic appearance of the stromata.

### Type

#### CHINA

Yunnan Province, Kunming, Luquan Yi and Miao Autonomous County, Tuanjie Town, Damaidi, causing root rot of *Quercus variabilis* Blume (*Fagaceae*), 5 August 2024, Jing Song, (**HKAS148941, holotype**), ex-type culture GBWSCC202508860, HKAS151626 (dried culture); *ibid* 6 September 2024, Senanayake IC (**HKAS148942, Paratype**).

### Description

Form from roots of dying host plants and then arise from rhizosphere soil. Teleomorph: Stromata upright, slender to plump fleshy, 20–50 cm total height, with fertile parts 10–20 cm high, 7–10(–15) cm diam., 250–600 g weight, variable size and shape, ranging from cylindrical, fusiform, turbinate to palm-shaped, terete to flattened, simple to branched from the base, or at the top, arising from long rooting stipes, with flattened sterile apices. Outer crust 200–300 μm thick, coriaceous, black, ostiole openings appear as black dots, ostiolar papillae distributes in this layer. Aerial part fertile, surface covered with a long persistent, powdery, whitish yellow fibrous substance, whitish in young stromata, turning progressively dark yellow at maturity. Immersed part 20–30 cm high, 3–6 cm wide, sterile consisted with stipes, cylindrical, black, no powdery substance, smooth. Interior tissues solid, homogenous to fibrous, spongy, off-white to yellow, with a slightly darker core in aged specimens, discolouration absence with 5% KOH. Ascomata 400–750 μm diam., 1200–1500 μm high, perithecial, aggregated, scattered, immersed in stromata, subglobose, coriaceous, central papillate, ostiolate. Ostioles 150–280 μm diam., raised-discoid, black, with a cylindrical papilla at center. Paraphyses 4–7 μm broad (x̄ = 5 μm), sparse, hyphal, thin-walled, narrowing towards the ends, hyaline, septate, slightly constricted at the septa, unbranched, longer than asci, embedded in mucilaginous matrix. Peridium 70–100 µm (x̄ = 80 μm), thick, dark brown, turning subhyaline inwardly, contain cells of *textura porrecta* mixed with *textura angularis*, evenly ticked. Asci (6–)8-spored, unitunicate, cylindrical, long-stipitate, 115–160 × 8–10 µm (x̄ = 2.5 × 3 μm) total length, the spore bearing parts 90–100 × 8–10 µm, with bilobed, apical apparatus, 2–3 µm high, 3–5 µm wide (x̄ = 3 × 4 μm) in water; cuboid, amyloid, 2–3 µm high, 2.8–3.2 µm wide (x̄ = 2.5 × 3 μm), in Melzer’s reagent; inverted trapezoid, reddish brown in Lugol’s solution, 2.5–3 µm high, 3–4.5 µm wide (x̄ = 3 × 4 μm), turning blue in Lugol’s solution after treat with 3% KOH. Ascospores 12–17 × 5–7 µm, overlapping uniseriate in the ascus, globose to oval or fusiform with acute ends, hyaline when young, becoming brown at maturity, with single, central guttule or with a large central and two, small polar guttules, smooth, thick-walled, with a conspicuous, S-shaped germ slit one-half to four-fifths spore length on the flattened side. Anamorph: nodulisporium-like anamorph with periconiella-like branching patterns. Mycelium consisting of septate, branched, smooth-walled hyphae, hyaline to pale brown. Conidiophores 17–25 µm length, 6–8 µm width (x̄ = 20 × 7 µm, n = 10), macronematous, mononematous, erect, septate, smooth, pale to dark brown, arising singly from hyphae; branching profuse and arborescent (periconiella-like), with primary and secondary branches producing numerous short terminal branchlets. Conidiogenous cells 7–10 µm length, 3–6 µm width (x̄ = 9 × 5 µm, n = 20), terminal, polyblastic, sympodially proliferating (nodulisporium-like), denticulate; conidiogenous loci minute, with small truncate scars. Conidia 6–9 µm length, 3–5 µm width (x̄ = 8 × 4 µm, n = 25), produced singly on denticles, ellipsoid to ovoid, hyaline, smooth, unicellular.

### Culture characteristics

Colonies after 7 days incubated at 20 °C in dark; on PDA reaching to 5 cm diam., circular, smooth margin, flat, bearing abundant aerial mycelium, initially white turn to yellowish-white with time, lack of pigment production, heavily sporulate with black, globose conidiomata; on MEA reach to 4.5 cm, circular, thin, arachnoid, with abundant, initially white, later turn to blackish brown, radiating, dendritic hyphae; center compact, slightly olivaceous to greyish; margin even to diffuse, reverse view blackish, radiating, dendritic hyphae start from the center spread towards the margin, lack of pigment production, fairly sporulated with black, globose conidiomata; on OMA reach to 5 cm, circular, white, dense, cottony to floccose, with abundant aerial mycelium, center compact and greyish, surface faintly radially striate with weak zonation, margin entire to diffuse, reverse view with circular, dark olivaceous to olive-brown pigmentation; central region buff to cream, sporulation not occurred.

### Additional material examined

#### CHINA

Yunnan Province, Xishuangbanna Dai Autonomous Prefecture, Mengla County, Mengla Town, 14 July 2025, Xiao Aka, (HKAS150824); *ibid*, Chuxiong Yi Autonomous Prefecture, Lufeng City, Yipinglang Town, 10 July 2025, Xie Tao, (HKAS150823).

### Notes

In the combined multilocus phylogeny analysis of ITS, LSU, *TUB2*, and *RPB2*, the collections of *Dianjunus* species clustered together forming a distinct clade with support value of ML/BI=100%/1.00 (Fig. 1). The bootstrap value in phylogenomic analyses also >90% (Fig. 3). All the collections of *Dianjunus rex* are morphologically similar, and stromata of tropical collections are larger than sub-tropical collections (Fig. 4). The young stromata are mostly with bulb-like apex and they form fingers-like branches when mature. The fresh stromata are very fleshy and dehydrate rapidly after detached. There is no strong smell or taste from stromata. Therefore, based on morphological characters and phylogeny, these collections are introduced as a new species, *Dianjunus rex*.

Our collections are morphologically resembled to genus *Squamotubera* P. Henn. Hennings (1903) erected the genus *Squamotubera* for the single species, *S. le-ratii* P. Henn., which was collected from Noumea, New Caledonia (island between Australia and New Zealand) from soil. Lloyd (1917) noted that *Squamotubera le-ratii* is similar to *Sarcoxylon compunctum* (Jungh.) Cooke except its horizontal or upright stromatal development from bark of living plants. Therefore, it was synonymized as *Sarcoxylon le-ratii* (P. Henn.) Lloyd. Rogers (1981) also argued that *Squamotubera* might congeneric with *Sarcoxylon.* However, considering the upright stromatal development of *Squamotubera* from subterranean sclerotia, Rogers (1981) conserved *Squamotubera* as a distinct genus and mentioned the requirement of further studies to clarify it. The genus *Squamotubera* is only known from its short Latin prologue by Hennings (1903) with no illustrations. Rogers (1981) noted that the holotype specimen of *Squamotubera le-ratii* is not available in specimen repositories (Farlow Herbarium, FH).

In the original description, Hennings (1903) described the stromata of *Squamotubera le-ratii* as “*tuberiform, round depressed with squamose external surface*”. Rogers (1981) amended the description and mentioned, “*the suspicion that stromata arise from subterranean sclerotia*” and “*the exterior is convoluted with smaller wrinkles overall*”. However, our collections do not fit any of these two characteristics. It is not “*tuberiform”* neither “*squamose or with wrinkles overall*”. Apart from these diagnostic characteristics, there are other important taxonomic traits that does not fit with the description provided by Hennings (1903) or Rogers (1981) such as “*interior pallid subcarnose*” or “*tan, solid with irregular lacunae*”; while in our fungus, the inner context was “*spongy, off-white to mostly yellow*”.

Rogers (1981) described, “*Capitate apices quickly become soft with gelatinous areas beneath the perithecial layer*” a characteristic never recorded in our specimens, whose context always remain fibrous, spongy. Additionally, Rogers (1981) described “Parasite sur les vielles souches de *Casuarina*” in French, which means “*Parasite on the old stumps of Casuarina sp.*”, a host plant not recorded for our fungus. Germ-slits in ascospores were not reported from *S. le-ratii* by Hennings (1903) and Rogers (1981) while our collections consist with S-shaped germ-slits. The stromata of *Squamotubera* are 7 cm height, 5 cm width while stromata in our collections sized with 20–50 cm height and 7–10 (−15) cm diam. Therefore, our collections are significantly different from *Squamotubera le-ratii* (Table 4).

**Table 4.**
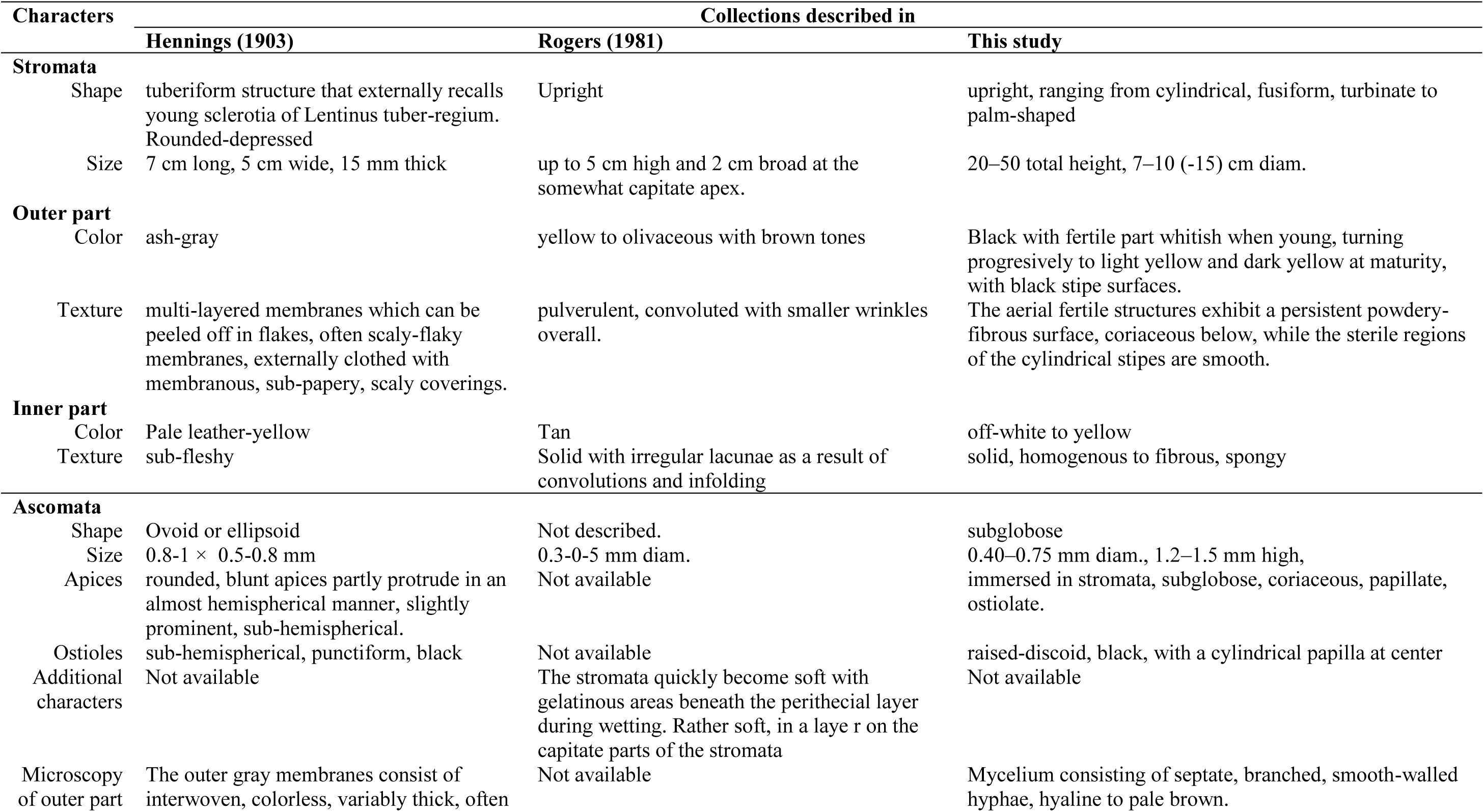
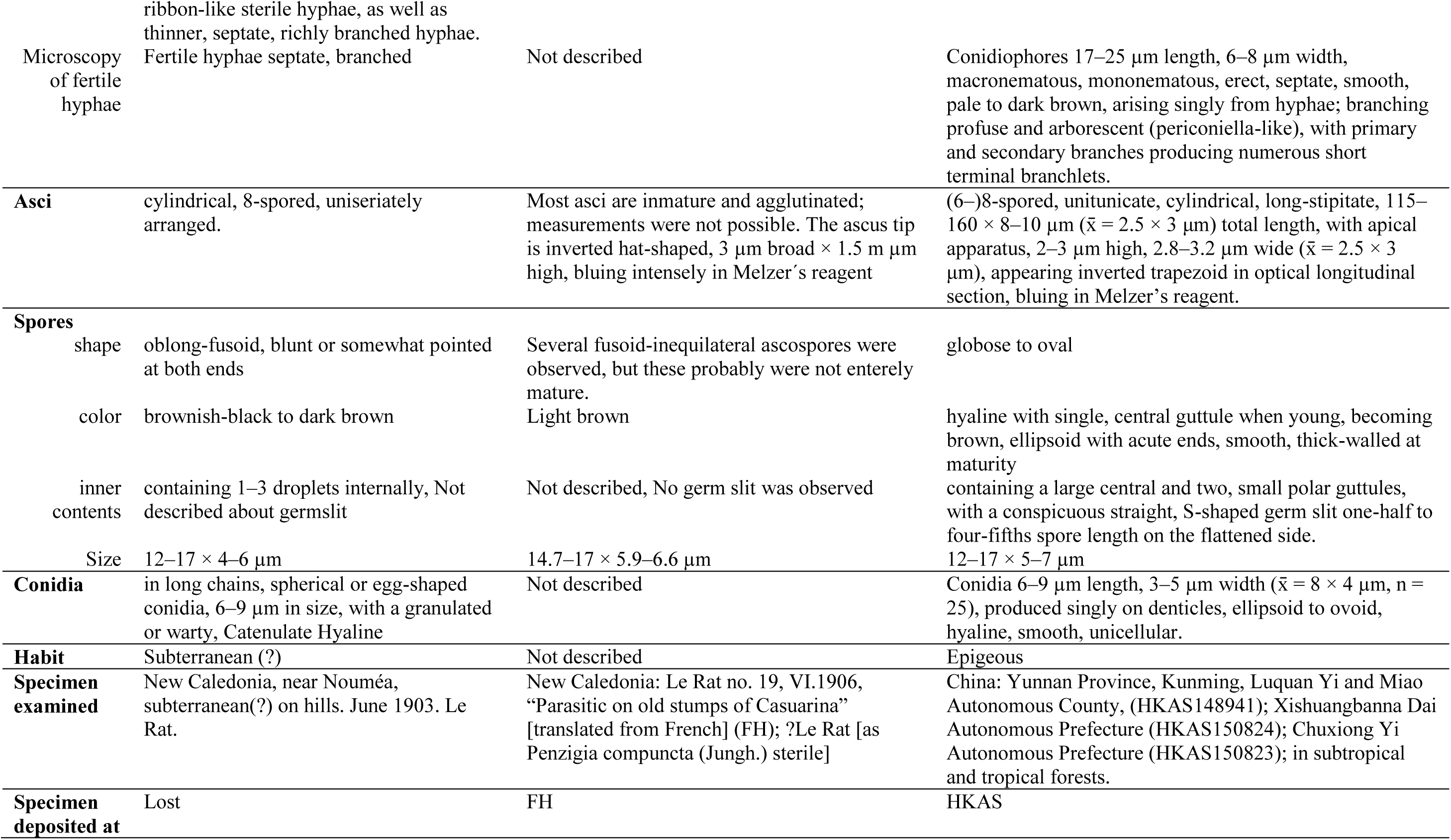
Morphological characters comparisson of collections of *Squamotubera* and *Dianjunus*.

## Discussion

Yunnan Province, situated in south-western China, is recognized for its exceptional fungal diversity (Feng and Yang 2018). Its varied topography, encompassing subtropical forests, alpine meadows, grasslands and wetlands, creates a spectrum of ecological niches and microhabitats that support a rich mycota (Sun et al. 2017). Driven by complex landscapes, diverse floristic communities hosting numerous endemic species, and predominantly tropical to subtropical climates, Yunnan constitutes a biodiversity hotspot for fungi. While over 6,000 species have been documented, total fungal richness is estimated to potentially exceed 100,000 species, with frequent discoveries of novel taxa and genera continuing to highlight the regiońs underexplores status (Senanayake et al. 2022). Research has extensively documented major phyla such as *Ascomycota* and *Basidiomycota*.

The Province hosts a significant diversity of ectomycorrhizal fungi, which form symbiotic associations with tree roots, particularly in temperate and subtropical forests (Liu et al. 2009). Commonly encountered genera include *Boletus*, *Ganoderma, Russula*, *Tricholoma*, *Tuber* (truffles) and *Xylaria.* These fungi play critical roles in nutrient cycling and forest ecosystem functioning. Yunnan is also notable for its edible fungi, especially *Morchella* species and truffles (genus *Tuber*), which hold substantially culinary and economic value both locally and globally. The utilization of wild edible mushrooms represents an important economic activity though it raises concurrent concerns regarding sustainable harvest and ecological impact. Thus, the diversity of edible fungi is a major focus of both ecological and economic research.

Up to now, very few species in *Xylariales* has been reported with production of massive stromata. The size of the stromata in *Xylariales* depends on the species, mode of life and geographical location of collections. *Engleromyces goetzei* P.Henn. is the ever reported, massive sized species in *Xylariales*. *Engleromyces goetzei* produces subglobose stromata seated on and partially enveloping bamboo culms forming two lobes (Whalley et al. 2010). Gibson and Kimeria (1962) reported *Engleromyces goetzei* from North Kenya, where it forms subglobose stromata growing towards the tops of on bamboo stems, about 30–40 ft. from the ground, with considerable variable size from small growths weighing a few ounces to the largest with a diameter of nearly 30 cm and weighing around 3.2 kg. Meanwhile, *Engleromyces sinensis* M.A. Whalley, A. Khalil, T.Z. Wei, Y.J. Yao & Whalley sampled in Yulong County in Yunnan Province, China presented much smaller stromata enveloping bamboo culms forming two lobes, globose to subglobose, with sizes of 4.3–4.9 × 4–5.5 cm width and 1.6–4 cm in height (Whalley et al. 2010). Most other morphologically resemble species with *Dianjunus rex* in *Xylariales* are *Daldinia concentrica* (Bolton) Cesati & de Notaris, *Xylaria hypoxylon* (L.) Grev., *X. cubensis* (Mont.) Fr., and *X. longipes* Nitschke and however, most of them are produce black stromata. However, *Dianjunus rex* differentiates, apart from their bigger size, by less branched, pale to dark yellow stromata with blunt apex (Phillips.et al. 2005).

Beyond its morphological distinctiveness, *Dianjunus* rex is phylogenetically isolated from all other known taxa in *Xylariales*. Divergence time estimates by Samarakoon et al. (2022) place the split between *Amphisphaeriales* and *Xylariales* at 154 (117–190) million years ago (MYA), with crown ages of 127 (92–165) MYA and 147 (111–184) MYA, respectively. Our analysis estimates the family *Dianjunaceae* diverged at approximately 65 (50–80) MYA (Fig. 3), a timeframe consistent with the common family-level divergence trend of 50–150 MYA reported for fungi (Hyde et al. 2017). The phylogenomic topologies generated by the concatenation and summary-coalescent (ASTRAL) approaches were largely congruent, with one notable exception in a non-target lineage. Specifically, the phylogenetic position of *Stanjemonium grisellum* exhibited topological discordance between the concatenation tree and the two ASTRAL species trees (which were perfectly consistent with each other) (Suplimentary data). In the ASTRAL analyses, the local posterior probability (LPP) for this discordant node was 0.84 for the BUSCO dataset and dropped to 0.62 for the OrthoFinder dataset (See suplimentary data). This moderate-to-low support indicates historical gene tree conflicts, likely arising from widespread incomplete lineage sorting (ILS) or historical hybridization events within this non-target group. Crucially, despite this isolated background conflict, our core target topology was flawlessly and robustly preserved across all three distinct workflows. *Dianjunus* was consistently resolved as a sister group to the *Graphostromataceae* with maximum support (Bootstrap = 100 in the concatenation tree; LPP = 1.0 in both ASTRAL trees). This definitive consistency across diverse datasets and evolutionary models demonstrates that the taxonomic and evolutionary placement of our target genus is unequivocally stable.

## Conclusion

This study elucidates the taxonomic identity, and phylogenetic relationships of an exceptional macromycete discovered in the subtropical and tropical forests of Yunnan Province, China. Through an integrative taxonomic approach combining detailed morphological examination, multi-locus phylogenetic analysis (ITS, LSU, *RPB2*, *TUB2*), and phylogenomic data, the fungus is formally described as *Dianjunus rex* gen. et sp. nov. Phylogenetic reconstructions robustly place this novel taxon within the order *Xylariales*, where it forms a distinct, well-supported lineage sister to the family *Graphostromataceae*. This phylogenetic isolation, coupled with a unique suite of morphological characters, including massive, upright, fleshy stromata, a persistent powdery-fibrous surface, and a nodulisporium-like anamorph, is proposed to establish the new family *Dianjunaceae*. Divergence time estimation, calibrated using a genome-scale dataset, indicates that the *Dianjunaceae* lineage diverged during the early Paleocene, approximately 65 (50–80) million years ago. This timing coincides with the Cretaceous–Paleogene (K–Pg) boundary and the subsequent period of significant planetary ecological reorganization. The emergence of this morphologically distinct lineage in the aftermath of a major extinction event may reflect an adaptive radiation or the exploitation of newly available ecological niches during forest ecosystem recovery. The stromata of *D. rex*, reaching dimensions of up to 50 cm in height and 2.2 kg in weight, represent the largest stromata documented within the *Ascomycota*. This discovery significantly expands the known morphological disparity of the *Xylariales*, demonstrating that the evolution of gigantism in ascomycete fructifications has occurred independently in multiple lineages, including the stromata-forming *Xylariales* and the apothecia- or truffle-forming *Pezizales*. The discovery of *Dianjunus rex* and the *Dianjunaceae* underscores the profound and largely unexplored fungal diversity within East Asian forests, particularly in the biodiverse regions of southwestern China. It affirms the critical importance of continued mycological exploration integrated with modern molecular systematics, which remains essential for revealing hidden lineages, understanding the evolutionary trajectories of fungal morphology, and elucidating the complex history of Earth’s ecosystems.

## Acknowledgement

This research was funded by the National Natural Science Foundation of China (Grant No. W2532026), the Yunnan Technology Innovation Program (202205AD160036), and the Yunnan Revitalization High-end Foreign Talent Support Program to Indunil C. Senanayake and Jesús Pérez-Moreno.

## Ethical statement

No ethical statement was reported.

## Adherence to national and international regulations

All the fungal strains used in this study have been legally obtained, respecting the Convention on Biological Diversity (Rio Convention).

## Author contributions

Formal Analysis (J-S, F-Z, Z-Y, C-Y); Resources (J-S, T-X, I-CS); Writing - original draft (Z-Y); Project administration (F-Y); Writing - review and editing (J-P-M, I-C-S); Supervision (J-P-M, R-C); Data curation (S-Y); Visualization (L-S, D-D); Validation (D-D, J-L); Methodology (Y-W); Conceptualization (D-L, F-Y, W-L); Investigation (X-S); Software (S-W); Funding acquisition (I-C-S, F-Y); Project administration (F-Y). Jing Song and Zhuyue Yan contributed equally to this work and share first authorship. All authors have read and approved the final manuscript.

