## Supplementary material for "Unearthing a fungal giant: *Dianjunaceae* fam. nov., a novel Paleocene lineage of *Xylariales* harbouring *Dianjunus rex* gen. et sp. nov.": Suplimentory figure 1 and 2

**Coalescent-Based Species Tree Inference Using BUSCO Dataset**

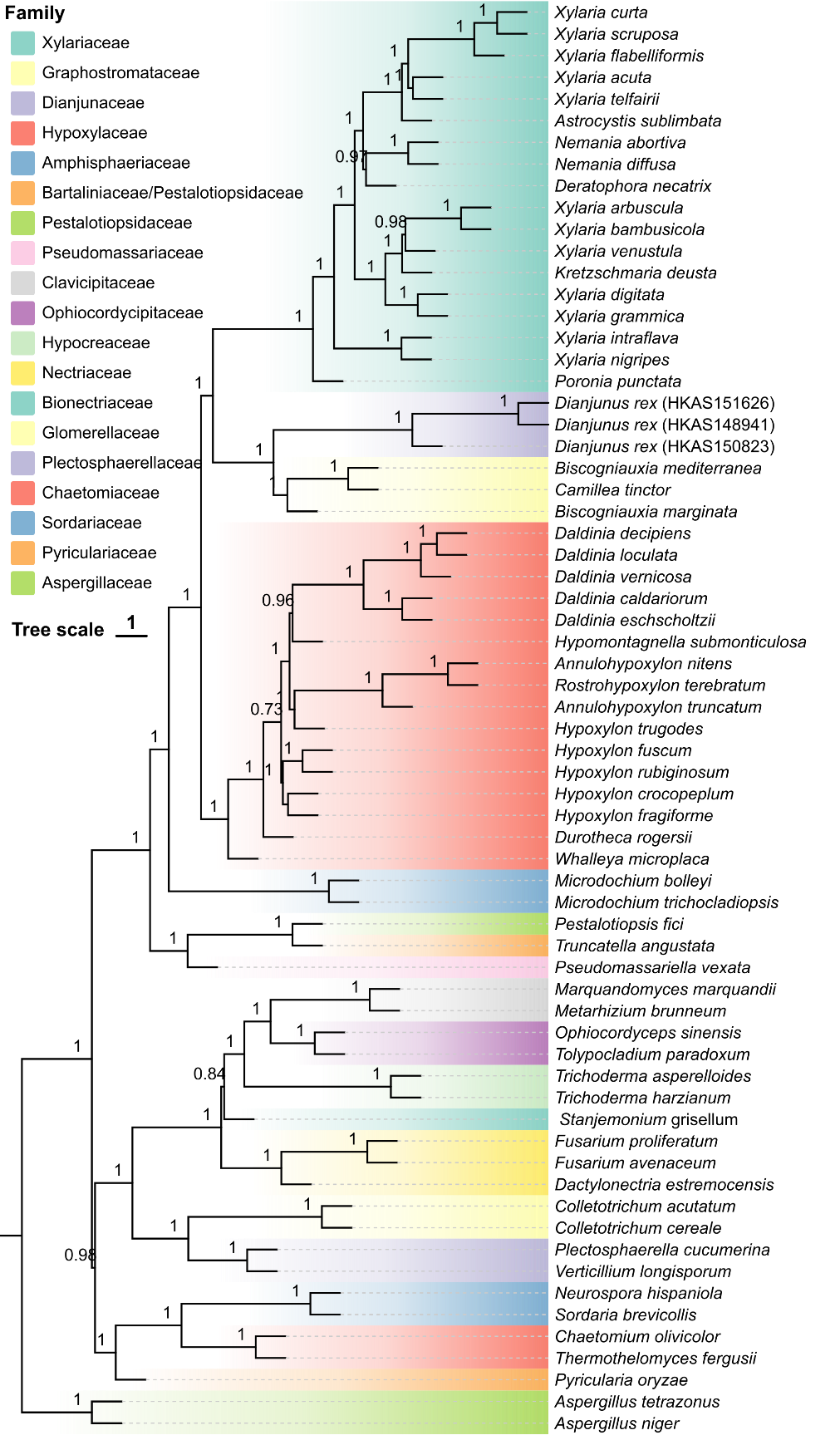

To evaluate potential gene tree conflicts and verify the robustness of the topology generated by the concatenation approach, a coalescent-based species tree was also reconstructed using the same BUSCO dataset. Individual maximum-likelihood gene trees for the 302 single-copy orthologs were inferred using IQ-TREE3 (v3.0.1) under the auto-selected best-fit evolutionary models (MFP) with 1,000 ultrafast bootstrap replicates, utilizing the same outgroup (*Aspergillus niger* and *A. tetrazonus*). The resulting gene trees were subsequently used as input for ASTRAL (v5.7.1) to infer the final species tree topology.

**Alternative Phylogenetic Inference Based on OrthoFinder Dataset**

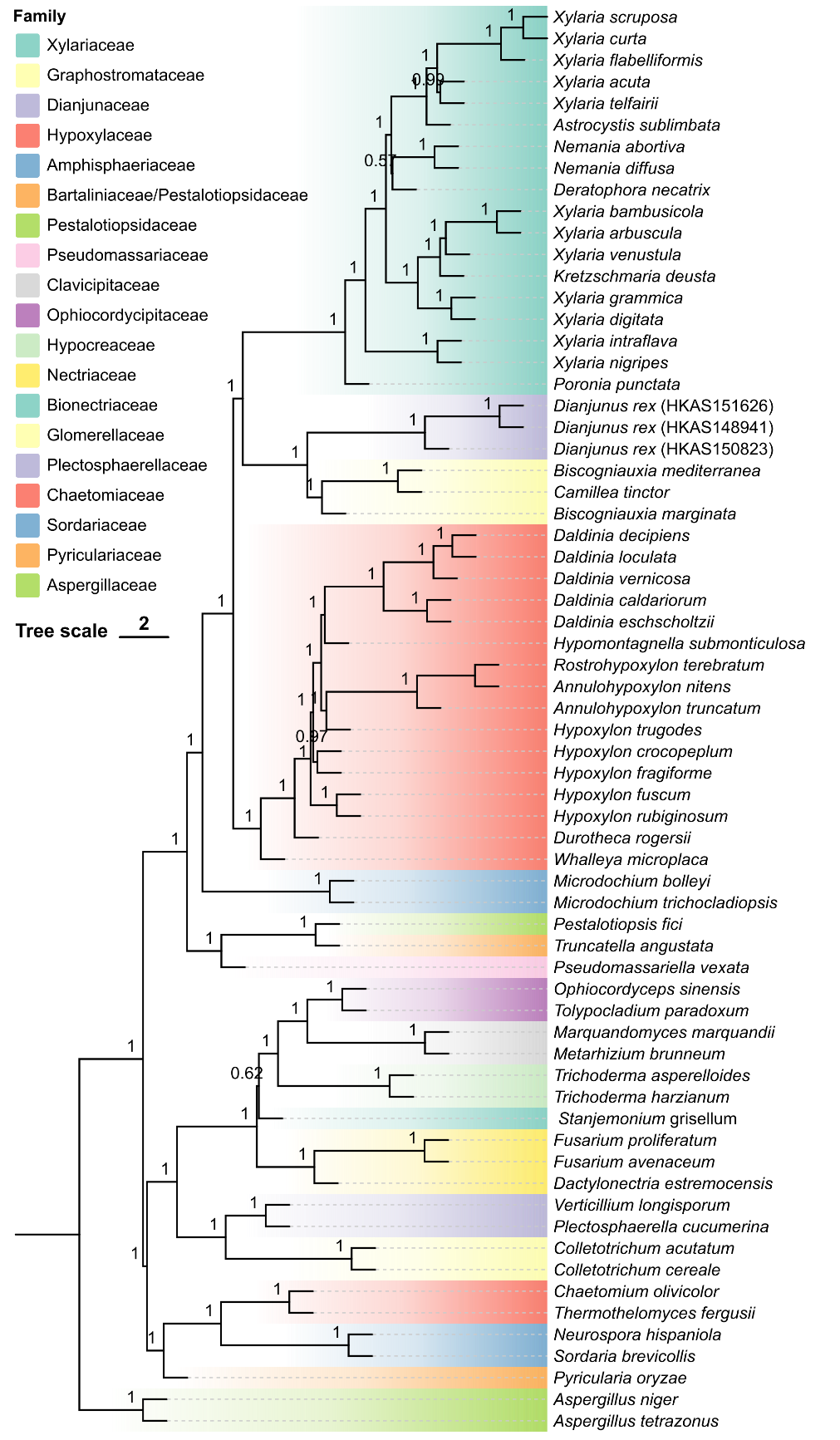
